# Cue-driven microbial cooperation and communication: evolving quorum sensing with honest signalling

**DOI:** 10.1101/2023.04.16.537056

**Authors:** Tamás Czárán, István Scheuring, István Zachar, Szabolcs Számadó

## Abstract

**Background:** Quorum sensing (QS) is the ability of microorganisms to assess local clonal density by measuring the extracellular concentration of signal molecules that they produce and excrete. QS is also the only known way of bacterial communication that supports the coordination of within-clone cooperative actions requiring a certain threshold density of cooperating cells. Cooperation aided by QS communication is sensitive to cheating in two different ways: *laggards* may benefit from not investing in cooperation but enjoying the benefit provided by their cooperating neighbors, whereas *Liars* explicitly promise cooperation but fail to do so, thereby convincing potential cooperating neighbors to help them, for almost free. Given this double vulnerability to cheats, it is not trivial why QS-supported cooperation is so widespread among prokaryotes.

**Results:** We investigated the evolutionary dynamics of QS in populations of cooperators for whom the QS signal is an inevitable side effect of producing the public good itself (cue-based QS). Using spatially explicit agent-based lattice simulations of QS-aided threshold cooperation (whereby cooperation is effective only above a critical cumulative level of contributions) and three different (analytical and numerical) approximations of the lattice model we explored the dynamics of QS-aided threshold cooperation under a feasible range of parameter values. We demonstrate three major advantages of cue-driven cooperation. First, laggards cannot wipe out cooperation under a wide range of reasonable environmental conditions, in spite of an unconstrained possibility to mutate to cheating; in fact, cooperators may even exclude laggards at high cooperation thresholds. Second, lying almost never pays off, if the signal is an inevitable byproduct (i.e., the cue) of cooperation; even very cheap fake signals are selected against. And thirdly, QS is most useful if local cooperator densities are the least predictable, i.e., if their lattice-wise mean is close to the cooperation threshold with a substantial variance.

**Conclusions:** Comparing the results of the four different modelling approaches indicates that cue-driven threshold cooperation may be a viable evolutionary strategy for microbes that cannot keep track of past behavior of their potential cooperating partners, in spatially viscous and in well-mixed environments alike.

## Background

Cooperation in the microbial world is abundant, mostly through excreted products benefiting not only the producer but other individuals, too. Microbial communities often rely on the production of metabolites or matrix substances that serve as common goods for the group as a whole. However, sacrificing valuable resources by the individual for the good of the group is a risky investment: cheaters may take advantage of honest cooperators by contributing less (or nil) to the common effort while they still enjoy the benefit. This means that selfish cheaters (“laggards”) have a growth advantage compared to cooperative producers, which, in the long run, leads to the tragedy of the commons (Hardin 1968) and the ultimate collapse of cooperation (Hamilton 1964, Diggle et al. 2007)).

In the face of the obvious fitness advantages of cheating in a cooperative group, it is puzzling how cooperation evolves and is maintained even in species with evolved mechanisms to avoid fraud. Aimed reward and punishment, straightforward antidotes to cheating, work only if group members can distinguish each other personally and remember the record of past actions of every group member back into a non-zero length of time. This is rarely the case even in vertebrate species, and much less in unicellular organisms with no brain or memory at all. Prokaryotes are therefore the least expected to harvest the benefits of cooperative group actions, lacking sophisticated mechanisms of partner recognition and record-keeping.

Yet there is an astonishing diversity and abundance of examples of genuine cooperation within and even between different prokaryotic strains producing public goods (West et al. 2006). These include the excretion of exoproducts like luciferin (Nealson et al. 1970), exoenzymes (Karray et al. 2018), bacteriocins (Ploeg 2005, Fontaine et al. 2007), siderophores (Stintzi et al. 1998, Popat et al. 2017), virulence factors (Rutherford and Bassler 2012) and biofilm matrix substances (Nadell et al. 2008), to mention just the most obvious forms of microbial cooperation. The actual functions of such different cooperative features may be connected in diverse combinations within the same strain, opening a wide range of complex microbial social strategies yet to be explored (Popat et al. 2017).

Most known forms of prokaryotic cooperation are threshold-limited: a certain number of nearby cooperators must all act simultaneously for the collective benefit of cooperation to exceed its individual costs (Chuang et al. 2010, Becker et al. 2018, Rosenthal et al. 2018). Thus, cooperating individuals have to coordinate their actions within a narrow spatiotemporal range, i.e., to synchronously express and excrete public goods in close proximity to one another.

Any mechanism ensuring that cooperators interact with other cooperators at a probability higher than their proportion within the population increases the chance of persistent cooperation. On the other hand, more frequent cooperator-cheater interactions increase the probability of cheater takeover. Since prokaryotic gene expression patterns are clonally passed down the generations, regardless of whether they are genetically or epigenetically determined, see e.g. (Veening et al. 2008)) with little room for phenotypic plasticity, the mechanism maintaining cooperation must always be a variant of kin selection. However, the actual form this mechanism takes may vary substantially.

The most trivial of such mechanisms is "environmental viscosity" (Strassmann et al. 2011). This means physical constraints limiting the mobility of individuals, thereby keeping the offspring adjacent to their parent and ensuring the overwhelming dominance of intraclonal interactions and effective kin selection thereof. It has been repeatedly shown that sufficiently high environmental viscosity can indeed maintain cooperation and prevent the invasion of cheaters in standard public good games (Nowak and May 1992, Kümmerli et al. 2009). Cooperators benefit from population viscosity in threshold public goods games as well (Vásárhelyi and Scheuring 2013), but they can stably coexist with cheaters even if the interaction size is limited and the population is perfectly mixed in this model context (Archetti and Scheuring 2010).

However, viscosity is rarely high enough in natural microbial habitats to effectively prevent cheater invasion from outside and/or constrain cheating mutants within. Therefore, in less viscous environments it may be of substantial selective advantage for the individuals to be able to size up the local density of potential cooperators and make actual cooperation dependent on that. This may prevent wasting valuable resources on futile attempts to cooperate locally when lacking sufficient cooperator density.

Quorum sensing (QS; (Miller and Bassler 2001)) is a simple genetic switching mechanism of communication-aided cooperation that may have evolved to provide this kind of phenotypic flexibility for unicells. It consists of a constitutively expressed signal, a membrane receptor, and an expression-excretion mechanism for cooperation (Figure 1). The QS switch triggers the transcription of certain genes upon sensing a sufficient number (a “quorum”) of cooperators in the neighborhood. The quorum is sensed by the capture of a sufficient number of signal molecules by a specific, dedicated membrane receptor, which then transmits the signal through an intracellular signal channel (often using cAMP) to the chromosome and activates the cooperation genes. Gram-positive bacteria use small autoinducing peptides as QS signals (Sturme et al. 2002), whereas Gram-negative bacteria (Schuster et al. 2013, Papenfort and Bassler 2016) and archaea (Paggi et al. 2003, Charlesworth et al. 2017) usually excrete N-acyl homoserine lactones for the same purpose. The fundamental signaling mechanism is the same in both cases. Quorum sensing has been discovered in almost any bacterial strain in which it was looked for, some strains utilizing multiple different QS systems (Henke and Bassler 2004), sometimes in synergy with each other. For example, the two principal QS systems (*las* and *rhl*) regulating the expression of virulence factors in *Pseudomonas aeruginosa* have been shown to form a reciprocal signalling network that synergistically enhances and tunes the strain’s reactivity to its physical and “social” environment (Thomas et al. 2023).

**Figure 1.**
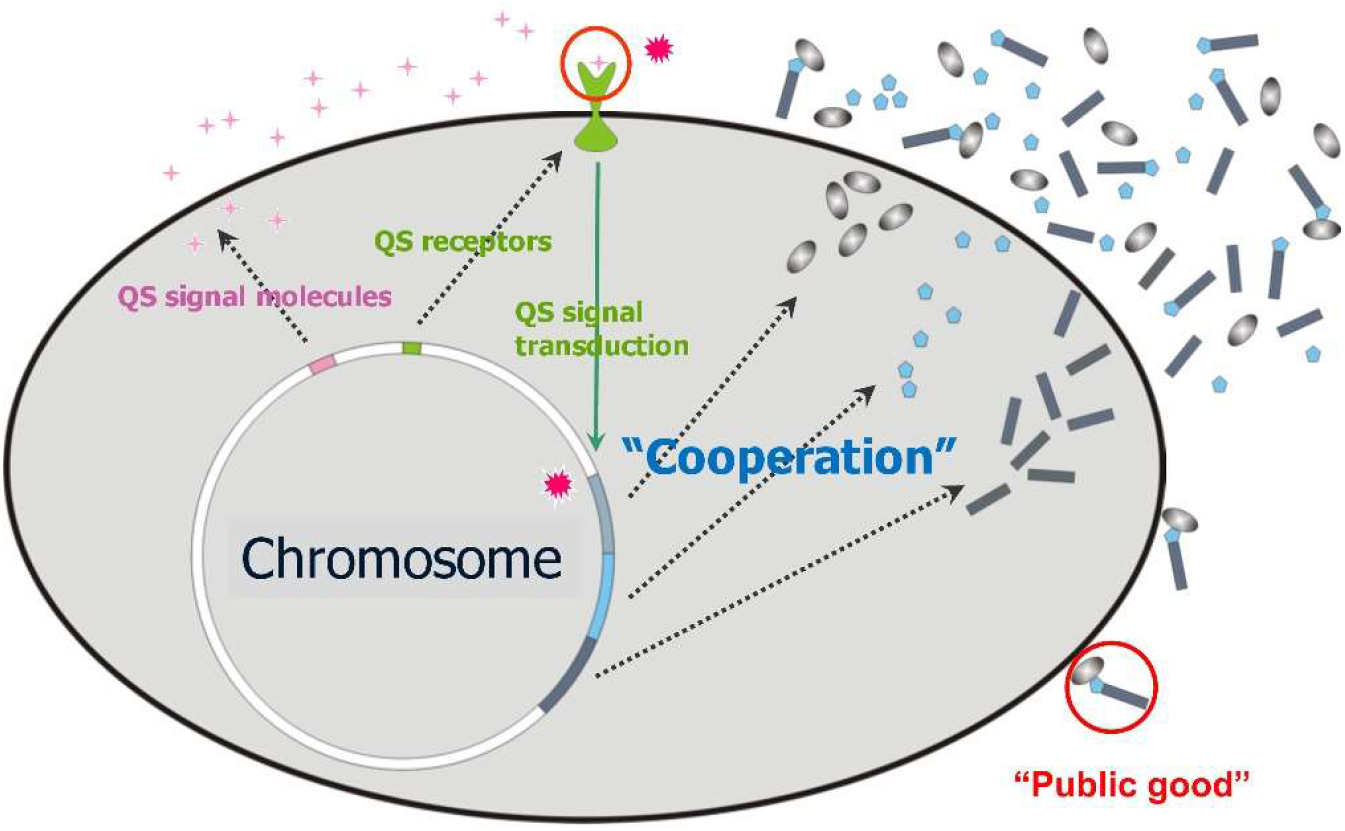
A simplified representation of quorum sensing regulated cooperation for the production of common goods collectively utilized by bacteria having access to them. The three basic genetic components of the QS-regulated cooperative system are a) the signaling component (pink), which comprises the genes for signal molecule production and excretion of signal molecules (stars); b) the signal detection and transduction system (green), including genes for the signal receptor and the second messenger system; c) the cooperation genes (blue) which are transcribed and expressed (and products externalized) if the extracellular concentration of signal molecules is sufficiently high. Red asterisks signify component events of signal detection: signal capture and signal transduction.

There are different interpretations of the function of quorum sensing (see (Cornforth et al. 2014)). The first interpretation is that it aims at sizing up the local density of potential cooperators (Fuqua et al. 1994, Salmond et al. 1995, Swift et al. 1996), whereas the second one is that it serves to assess the diffusibility of exoproducts in the environment (Redfield 2002). These two functions are difficult to disentangle as both high cell density and low diffusion can lead to a high local concentration of the QS signal. There is some indication that some bacterial strains might be able to distinguish these two types of information using combinatorial QS signals, i.e., by using two different signal molecules with covarying decay and autoinduction rates (Cornforth et al. 2014).

Either way, as QS is a communication system, it is prone to defection or deceit by another type of cheater: individuals with a silent set of cooperation genes but capable of sending out false signals of intent for cooperation (lying) may still enjoy the full benefit of the public good produced by nearby cooperators responding to the false signal. A textbook example of such defectors (“liars”) is the *lasR* mutant of *Pseudomonas aeruginosa* that does not respond to QS signals of the wild type and, consequently, does not cooperate in producing an important public good, a protease exoenzyme virulence factor (Diggle et al. 2007). The cheating mutant has been shown to enjoy a substantial reproductive advantage over cooperative strains (Chen et al. 2019) for the obvious reason of not carrying the metabolic burden of cooperation (exoenzyme production).

A more nuanced way of cheating is increasing signal production while evolving a higher threshold value for cooperation. Brown and Johnstone in a seminal model of QS were able to show that increasing conflict of interest (decreasing relatedness) favors such ‘coercive’ variants (Brown and Johnstone 2001). These can manipulate older strains with lower thresholds into increased production of the public good by mimicking a higher cooperator density with the increased signal production. In silico study of QS found that such coercive variants are more likely to emerge in genetically mixed populations with decreased relatedness (Wang et al. 2020).

If, however, the signal cannot be switched off, cheating is expected to be less deleterious for cooperation. For example, the signal may be the public good itself, as in the case of lactic acid bacteria producing the bacteriocin called nisin. Nisin production is QS-regulated (Kleerebezem 2014), but the QS signal is nisin itself, so bacteria producing nisin are also signaling. Cooperators always produce nisin at a low expression level, advertising their willingness to cooperate, and they express and excrete nisin at an elevated level when a quorum is reached. However, the cooperation signal can be faked by non-cooperators: the bacteriocin gene can be expressed constitutively at a low level to produce the signal, but cheaters never express it at sufficiently high levels to considerably contribute to the common good. Therefore, individuals capable only of low-level nisin expression are actually liars.

Another type of cheater is a cooperator that expresses more nisin in its signaling state than the normal signal level but less than the cooperation level. By issuing extra signals above the basic expression level of honest cooperators, these cheaters gain a fitness benefit by increasing the local signal concentration which, therefore, may reach the quorum threshold and convince more cooperators to join in the common effort, possibly in vain. This is equivalent to promising more cooperation than actually provided – a milder version of the liar strategy.

To study the dynamical and evolutionary properties of QS-regulated cooperation, it is sufficient to assume that the individuals clonally inherit three fundamental QS-related properties: 1) signaling (**S**) or not (**s**) the intention/ability to cooperate; 2) responding (**R**) or not (**r**) to above-quorum signal levels; and 3) cooperating (**C**) or not (**c**) when a quorum is reached. Each of these properties may be determined by a number of different genes, but the only trait we consider relevant is the functionality of the corresponding gene set as a unit. Therefore, we assume three loci with two functional alleles on each, which allows for 2^3^ = 8 different genotypes. For QS to hold, cooperation (i.e., the expression of the **C** allele) is assumed to be conditional on the presence of a critical number of signaler individuals (the quorum, i.e., those harboring **S** and/or **C**) within the interaction neighborhood of individuals possessing both **C** and **R**. In other words, cooperators capable of detecting the signal will cooperate only above the critical local quorum and mute their cooperation gene otherwise. Non-responder cooperators cooperate unconditionally, and they may issue the signal either at the normal expression level or at an elevated one.

The question we aim to answer is whether cue-driven (e.g., nisin type) cooperation and/or communication can be maintained in a population of quorum sensing microbes in the face of all possible mutations allowing cheater strategies, assuming different costs of cooperation, signaling, and signal detection/response. Here, we will scrutinize a family of models built on the above assumptions, allowing all three possible types of cheaters to appear. The models are built on the individual-based approach of Czárán and Hoekstra (Czárán and Hoekstra 2009), extending it both in scope and methodology of representation. For a deeper insight into the coexistence dynamics of the various strategies, we derive analytical 1) mean-field (MF) and 2) configuration-field (CF) (Czárán 1998) approximations besides the corresponding individual-based 3) non-spatial and 4) spatial stochastic simulations of the threshold public goods game (TPPG).

## Model basics

We used four different model types: MF approximation, CF approximation, non-spatial and spatial (on-lattice) agent-based models. They share the same basic assumptions (explained below), but gradually relax crucial simplifications while also losing analytical tractability. For the specifics of the different approaches, see Methods and models, for parameters, see Table 1.

**Table 1.** Model variables and parameters used throughout this study.

|  | Symbol | values in the configuration field model | values in spatial simulations |
| --- | --- | --- | --- |
| fitness of player $i$ | $w_i$ | Eq. 1 | Eq. 1 |
| fitness difference of players $i$ and $j$ | $\Delta w_{ij}$ | calculated from $w_i$ and $w_j$ | calculated from $w_i$ and $w_j$ |
| probability that the offspring of player $i$ replaces player $j$ | $p_{ij}$ | Eq. 2 | Eq. 2 |
| population size | $P$ | ∞ | 90.000 |
| neighborhood size | $N$ | 9 | 9 |
| selection strength | $\sigma$ | 1 | 1 |
| cost of signal detection | $r$ | 0.01, 0.20 | 0.01, 0.05, 0.10 |
| cost of signaling | $s$ | N/A | 0.01, 0.05, 0.10 |
| cost of cooperation | $c$ | 0.3 | 0.2, 0.3 |
| baseline cost | $c_0$ | 1 | 1 |
| benefit of cooperation | $b$ | 0.3, 0.5, 0.6, 0.65 | 0.5, 0.8 |
| mean no. of diffusion steps per generation | $D$ | N/A | 0.0, 0.1, ..., 1.0 |
| quorum signal threshold | $Q$ | $Q = \kappa$ | $Q = \kappa$ |
| cooperation threshold | $\kappa$ | 3 | 2, 3, ..., 6 |
| functional mutation rate | $\rho$ | 0 | $0, 10^{-4}$ |
| grid size | $M$ | N/A | $300 \times 300$ |
| generation count | $G$ | 10.000 | 10.000 |

### Strategy set

Individual behaviors (strategies) are determined by three "functional genes" (heritable traits possibly encoded by a number of genes each) which control cooperation (C), extra signaling (S), and signal detection and response (R) (see Figures 1, 2). Each of these genes can be in one of two states (i.e., they have two "alleles"): they may be active (denoted by bold capitals: **C, S, R**) or inactive (bold minuscules: **c, s, r**). Note that an active cooperation gene (**C**) provides two things in the model: (*i*) the public good, (*ii*) and a baseline (cost-free) signal level (hence ‘cue-based’ cooperation). Accordingly, there are eight possible strategies ("phenotypes"); see Figure 2. "Lazy" (*La*: **csr**) never issues or detects the quorum signal and does not cooperate. "Trusty" (*Tr*: **Csr**) cooperates unconditionally, as it does not communicate, neither gives nor listens to signals. "Bouncer" (*Bo*: **CSr**) is also an unconditional cooperator issuing extra signals but not listening to them. We assume that the extra signal expression doubles the signal level so that a *Bo* individual counts as two signallers in its interaction group. "Smart" (*Sm*: **CsR**) is a cooperator that detects the quorum signal and cooperates if the signal level in its immediate vicinity exceeds the quorum threshold. "Nerd" (*Ne*: **CSR**) is a quorum-sensitive cooperator that always produces an extra signal dose. "Liar" (*Li*: **cSr**) issues the quorum signal but never cooperates. "Curious liar" (*Cl*: **cSR**) acts like Liar, but also detects the signal. Finally, "Voyeur" (*Vo*: **csR**) only detects the signal and never cooperates.

**Figure 2.**
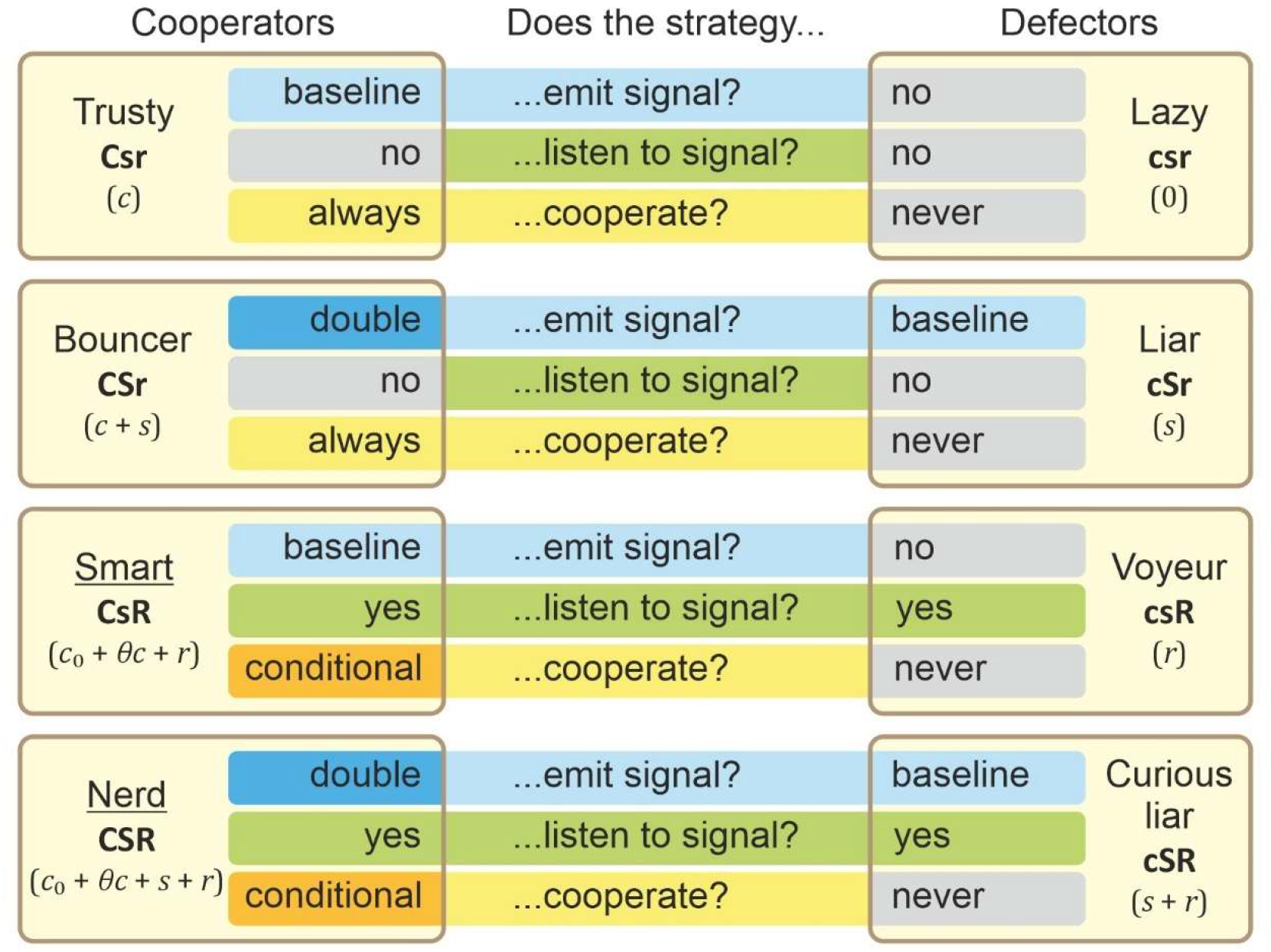
Strategy set of the QS model. Strategies are listed in boxes; their genotypes are denoted in bold typeface and their metabolic costs in parentheses. Capital letters in genotypes indicate expressed “genes”, minuscules indicate inactive alleles. *c*_0_ is the baseline metabolic cost paid by everyone, *c* is the cost of cooperation, *θ* = 1 if the quorum threshold is reached (otherwise *θ* = 0), *s* is the signal production cost, and *r* is the signal-detection cost. Underlined strategies are context dependent, capable of switching to cooperation when a signal quorum is reached. The individual strategies are characterized as follows: Lazy does nothing; Voyeur detects signal but does not cooperate; Liar signals but does not cooperate; Curious liar produces and detects signal but does not cooperate; Trusty does not communicate but always cooperates and thus also issues the QS signal; Smart detects the signal and cooperates if the signal level exceeds quorum; Bouncy issues extra signal but does not listen to it; Nerd produces the extra signal, detects the signal and cooperates if the signal level exceeds quorum

### Metabolic costs

Each player invests a fixed metabolic effort *c*_0_ into its own maintenance. This baseline metabolic burden is the same for all strategies. Cooperation (**C**), the emission of extra signal molecules (**S**) and the production, maintenance and operation of the signal response system (**R**) are all metabolically costly; the corresponding *c*, *s* and *r* costs are added to the baseline metabolic burden of the players expressing them, to yield the total metabolic cost of the corresponding genotype. The extra signal molecules and the signal response system are always expressed in all the players carrying the active genes for these functions. The expression of the cooperation gene **C** may be conditional on the concentration of nearby signal molecules (i.e., the number of signaling neighbors), provided that they express the signal response system **R** and thus they are capable of detecting the signal. Cooperating players also issue the cooperation cue at no additional cost. The condition for the expression of the cooperation gene in conditional cooperators (the Smart and the Nerd strategies) is that the number of signal doses within the QS neighborhood exceeds the QS threshold *Q*. Obviously, non-expressed cooperation genes carry no metabolic cost. Figure 2 summarizes the total metabolic costs of the strategies with the local signal levels (i.e., the number of signal doses within the QS neighborhood) below and above the quorum signal threshold.

### Cooperation benefit and fitness

The metabolic cost of a player with at least *κ* active cooperators in its own interaction group is reduced by a factor 0 < *b* < 1, which is the cooperation benefit. The benefit reduces the metabolic cost of the individual in a multiplicative manner. Notice that the cooperation threshold *κ* is not equivalent with the quorum signal threshold *Q* – these two thresholds are different, even if their values are the same numerically. *Q* is the minimum number of quorum signal doses within the interaction group of a conditional cooperator that is sufficient to switch its **C** gene on, whereas *κ* is the minimum number of active cooperators within a group necessary for members of the group to enjoy the cooperation benefit. We assume *κ* = *Q* throughout this study, implying that this relation is the evolutionary optimum for cooperators: any deviation by switching on cooperation too late or too early is selected against. The fitness *w*_i_ of a player *i* is linearly decreasing with its actual metabolic cost *C*_i_, that is:

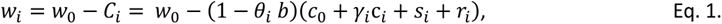

where *c*_i_ = *c* if the cooperation gene can be expressed in player *i*, otherwise *c*_i_ = 0; similarly, *s*_i_ and *r*_i_ are the corresponding costs of issuing an extra signal and responding to a quorum of signals by player *i*. *θ*_i_ = 1 if the number of active cooperators in the interacting group that *i* belongs to is at least *κ*, otherwise *θ*_i_ = 0, and *γ*_i_ = 1 if the number of signal doses (i.e., the number of cooperators plus the number of extra signalers) is at least *Q* within the group of *i*, or if *i* is an unconditional cooperator; *γ*_i_ = 0 otherwise.

### The metabolic cost of various strategies

Trusty always cooperates and signals, and thus it always pays the cost of cooperation (*c*), but it enjoys the cooperation benefit (*b*) only if the number of cooperators in its interacting group exceeds the critical threshold *κ*. Bouncer behaves as Trusty, except that it produces twice as many signal molecules as Trusty at some extra cost *s*. Smart cooperates (and pays the cost *c*) only if the number of signal doses is at least *Q* in its interacting group but operating the response system listening to the signal of the others implies a small cost *r*. Nerd is a conditional cooperator like Smart, producing an extra dose of signal at the extra cost *s*. The strategy Liar never cooperates and doesn’t listen to the quorum signal, but it does produce it, trying to induce conditional cooperators in its interacting group, again at the cost *s*. This model assumes that the quorum signal is the product of cooperation, thus the emission of a single dose of it is free of charge in all potential and actual cooperators, but they can produce an additional dose at a cost. Non-cooperators pay even for the first signal dose if they produce it, so that the Curious liar strategy both producing and detecting the signal pay the corresponding costs *s* and *r*; the Voyeur strategy pays only *r* for detecting the signal. Figure 2 summarizes the metabolic costs and benefits for all possible genotypes in interacting groups with the number of actual cooperators below and above the cooperation threshold *κ*.

### Reproduction, mutation, evolution

Reproduction takes place during pairwise interactions between individuals, following the rules of probabilistic imitation dynamics: player *i* has a chance of occupying the site of its opponent *j* with its own offspring. This has probability *p*_ij_, proportional to the relative fitness (cost advantage) of *i*, as 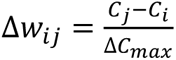, where Δ*C_max_* = *c* + *s* + *r* + *b c*_0_ the largest possible cost difference (between an individual expressing all functional genes but not receiving benefit, and another one expressing none of the genes but receiving the full benefit). Then:

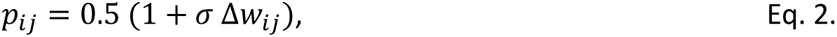

where *σ* is the strength of selection. Obviously, *p*_*ji*_ = 1 − *p*_*ij*_.

We assume that, during reproduction, any of the three functional loci (C, S and R) may mutate from its functional to its inactive form in the offspring, and back-mutations are also allowed, with each of the six possible mutation events occurring at its own specific rate (possibly zero). The dynamical equilibria and the trajectories of the resulting selection processes are the primary targets of this study, discussed in the four different modelling approaches.

## Results

### Mean-field (MF) model

In the MF model, we only consider three strategies (Lazy, Trusty, Smart), as in the limit, only these could be present in any equilibrium (see Methods and models and Appendix 1). It is easy to show that Lazy is stable against the invasion of the mutant strategies Trusty and Smart, if the sum of the initial frequencies of these mutants are below the threshold *K* − 1/*N* (where *κ*/*N* → *K*). Furthermore, it can be shown that the only alternative stationary state is the coexistence of the *La* and *Tr* strategies. This ensues if the benefit-to-cost ratio is sufficiently high (*b*/*c* > 1/(*c*_0_ + *c*)). However, this alternative fixed point is not stable against frequency fluctuations larger than 1/*N* (for more details, see Appendix 1). This result shows that QS communication does not convey any benefit in a well-mixed environment: cheaters will always be present, and unless the cooperation benefit exceeds its cost substantially, cooperators will be wiped out by cheaters; therefore, signaling and responding strategies have no chance to persist whatsoever.

### The configuration-field (CF) model

In the CF model we also considered only the three feasible strategies (Trusty, Smart and Lazy), because none of the others have a chance to persist in the presence of any of these three, just like in the MF approximation. The replicator dynamics yields four characteristically different outcomes:

- Case 1: If cooperation is too costly compared to the benefit for both Trusty and Smart, and Lazy is the only fixed point (stable one) of the system (background of Figure 3A).
- Case 2: With increasing benefit *b*, a stable and an unstable polymorphic state of Smart and Lazy emerge. Which one of the two stable fixed points (Smart/Lazy or Lazy) the system approaches depends on the initial frequencies (background of Figure 3B).
- Case 3: If the benefit of cooperation is high, but signal detection is costly, then Smart and Trusty swap roles: a pair of stable and unstable polymorphic Lazy/Trusty fixed points emerge besides the pure Lazy stable state, and the signal detection gene gets lost (background of Figure 3C).
- Case 4: Finally, when the signal detection cost is low, and the cooperation benefit is high enough, the system behaves like in Case 2 except that an additional unstable fixed point and a saddle point occur on the Lazy/Trusty polymorphic margin of the state space (background of Figure 3D). This, however, does not change the final state of the system compared to Case 2: it is either monomorphic Lazy or polymorphic Smart*/*Lazy, depending on initial conditions.

**Figure 3.**
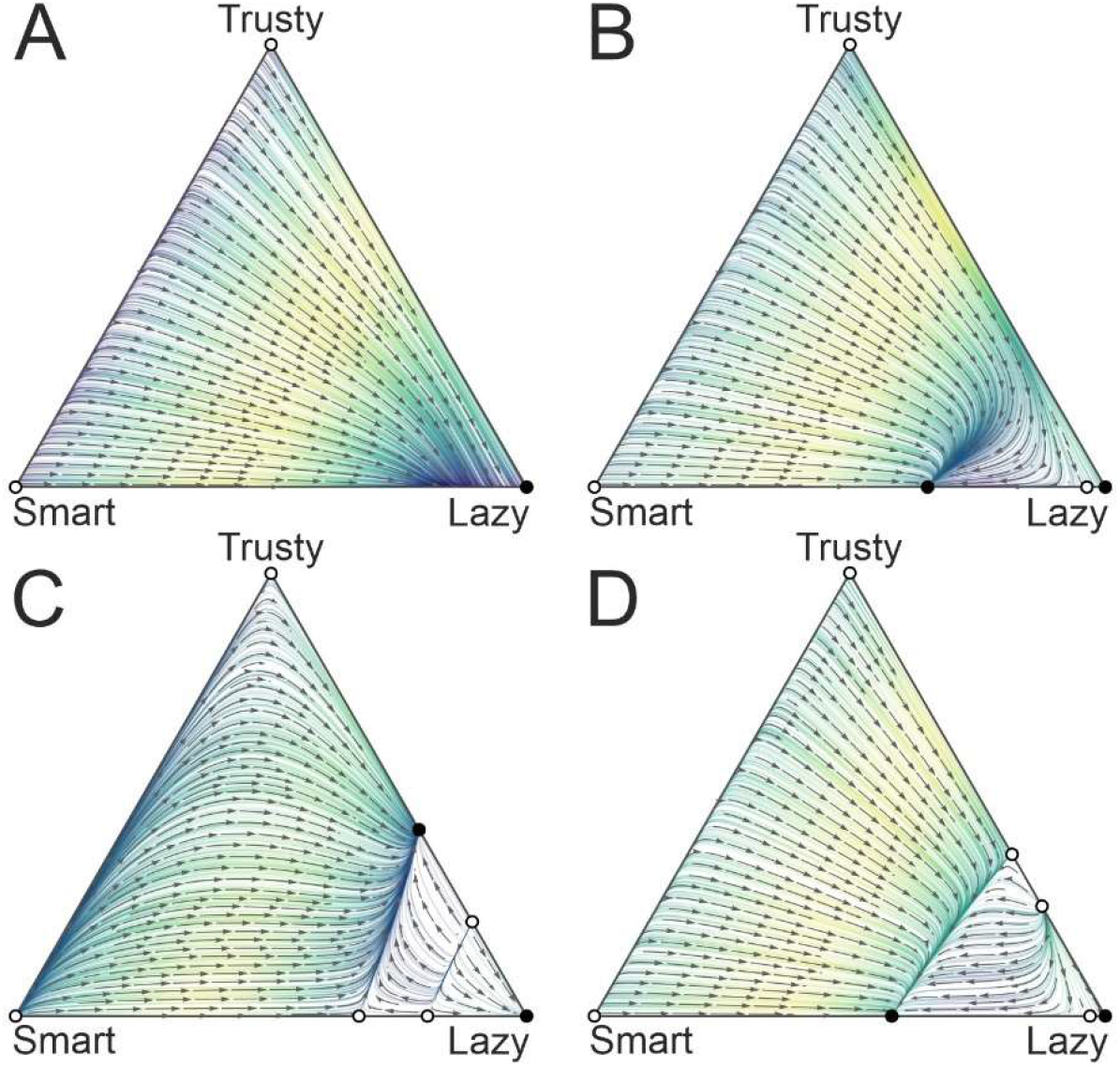
Configuration-field approximation, with the vector fields of the dynamics of the three feasible strategies (Lazy, trusty, Smart) in four different cases. **A**: Lazy is the only fixed point of the dynamics; parameters are *b* = 0.3, *r* = 0.01. **B**: The monomorphic Lazy and the polymorphic Smart/Lazy states are the stable fixed points of the dynamics; *b* = 0.5, *r* = 0.01, **C**: The monomorphic Lazy and the polymorphic Trusty/Lazy states are the stable fixed points of the dynamics; *b* = 0.65, *r* = 0.2, **D**: The monomorphic Lazy and the polymorphic Smart/Lazy states are the stable fixed points of the dynamics; *b* = 0.6, *r* = 0.01. All CF systems admit multiple unstable fixed points as well, including the Smart and the Trusty corners of the state space. In all cases, the rest of the parameters are *N* = 9, *κ* = 3, *c*_0_ = 1, *c* = 0.3.

For the detailed mathematical derivations of these results, see Appendix 2.

### Agent-based non-spatial model

To address the effect of the finite size of interacting groups, along with the spatial constraints arising from limited agent mobility and local (i.e., neighborhood-) interactions, we developed an agent-based implementation of the CF model, with its basic assumptions as presented above.

Not surprisingly, the trajectories of the non-spatial agent-based simulation model (Figure 34, row 3 red lines) closely trace the vector fields of the CF approximation (Figure 34 row 3 backgrounds, where the parameters of the cases A-D corresponds to Figure 3 A-D parameters) at any parameter setting. The only difference between the assumptions of the two models is that the CF approximation assumes an infinite population size, whereas, in the non-spatial simulations, lattice size is *P* = 90.000. This accounts for the stochastic noise on the simulated trajectories compared to the background CF vectors.

### Agent-based spatial (lattice) model

#### 3-strategy model without functional mutations

For comparative purposes, first we followed the trajectories of the three feasible strategies (Lazy, Trusty, Smart) in the lattice model, once again omitting those with no chance to persist in the MF and the CF approximations (see Appendix 1, 2) (Figure 4, rows 1-2). Mutations in any of the three functional loci were ignored. Results differ from those of the lattice-based CF approximation (Figure 4, trajectories of row 3, with the vector-field of the analytic CF as background), both at high and zero cell mobility (*D* = 10.0 and *D* = 0.0, respectively, see Figure 4, rows 1-2). The empirical (simulated) vector fields reveal that the spatially explicit model results in the least cooperative steady-state populations (backgrounds of Figure 4, rows 1-2, columns B-D): almost all the fixed points are at, or very close to, the Lazy corner of the strategy simplex. This means far worse conditions for cooperators compared to the CF approximation (Figure 4, row 3, columns B-D) that allows for polymorphic steady states at a considerable part of its parameter space. As the only difference between the CF and the well-mixed lattice model is that in the lattice model, the competing agents are immediate neighbors with overlapping cooperation neighborhoods, this may seem a counterintuitive outcome in view of the common understanding that spatial constraints like localized interactions, in general, help cooperators in two-person public goods games (Nowak and May 1992). The explanation for this peculiar result lies in the balance of two counteracting effects, each attributable to one of the two (convoluted) phases of the threshold cooperation game: 1) the fitness-acquiring cooperation phase, and 2) the competitive imitation phase.

1. In terms of fitness gains, cooperators do better in spatially “viscous” populations (spatially explicit case with *D* = 0.0, at which only very limited mixing occurs due to the copy of the winner being placed one site removed from its parent). When cooperators cluster together, it is mostly them who harvest the synergistic benefits of their own cooperative investment (in accordance with the conventional kin selection argument; see Figure 5A). Viscosity (low diffusion, *D* = 0.0) yields a polymorphic steady state on the Lazy-Smart boundary of the simplex compared to the non-viscous case (*D* = 10.0); compare rows 1 and 2 of Figure 4D (when the benefit is high and signal detection is cheap: *b* = 0.6, *r* = 0.01). Increasing agent mobility (high diffusion) approximates random spatial patterns in the limit, which is obviously detrimental for cooperators either because of the wasted cost *c* of cooperation if the cooperation threshold is not met in their neighborhoods, or due to their exploitation by parasitic free-riders if it is. In either case, parasites are better off in terms of fitness collected, and they prevail. That is, in the fitness-acquiring phase mixing is advantageous for parasites, whereas spatial correlations arising from limited mobility (viscosity) help cooperators to persist.
2. The outcome of the competitive imitation step of the TPPG between the members of cooperator-parasite pairs depends on the fitness collected by the players during the cooperation phase. In the spatially explicit model with strong mixing, the expected number of cooperators *E*(*n*_c_) in the two overlapping neighborhoods of a neighboring cooperator-parasite pair are expected to be the same for any cooperating (C: Trusty or Smart) individual playing against a neighboring parasite (P: Lazy): *E*_c_(*n*_c_) = *E*_p_(*n*_c_). This follows from the facts that a) adjacent individuals are members of each other’s neighborhoods; b) the members of the remaining 7-7 individuals in their neighborhoods are either exactly the same (in their overlapping parts) or c) they are (statistically) equivalent due to the intensive mixing assumed (cf. Fig. 5B). Therefore, the expected cooperation benefit is the same for the two interacting players, but the parasite spares the cost of cooperation and/or signal detection, enjoying thus a fitness advantage over any cooperator. In the CF model, a) the two interacting players are not neighbors; therefore, b) their cooperation neighborhoods do not overlap, but c) the remaining 8-8 individuals in their two independent neighborhoods are again identical in the statistical sense. Thus, the expected number of cooperators in the cooperating player’s neighborhood is always larger by 1 than in the neighborhood of its parasitic opponent: *E*_c_(*n*_c_) = *E*_p_(*n*_c_) + 1, due to the focal cell being a cooperator (Figure 5B). This difference explains the relative disadvantage of cooperators in the lattice model with intensive mixing compared to the CF model (as demonstrated between rows 2 and 3 of Figure 4).

**Figure 4.**
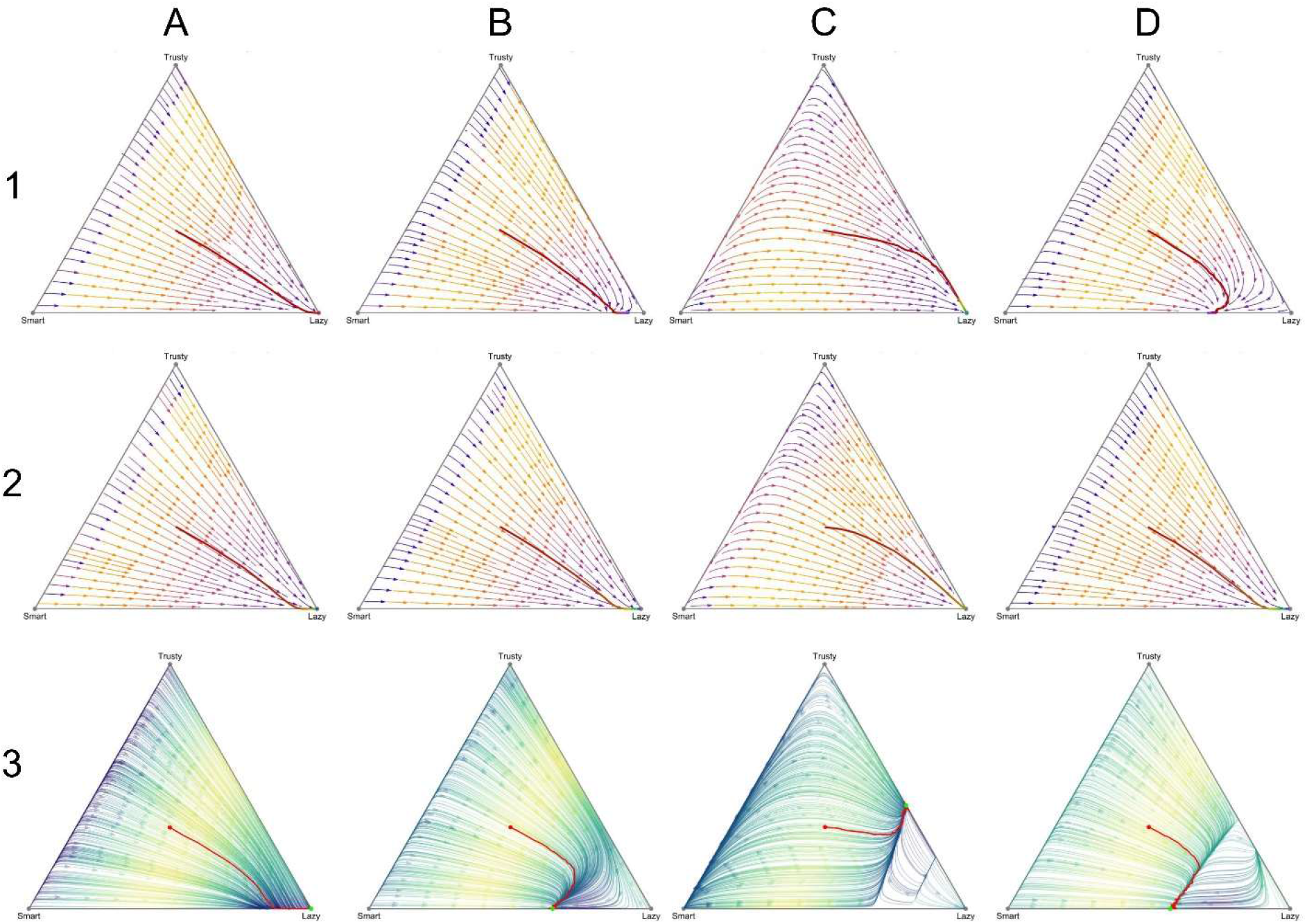
Sample trajectories of the configuration-field (CF) and the spatially explicit lattice model on the corresponding vector fields. Functional mutations have not been considered here. **Row 1**: Spatially explicit simulations, no mixing (*D* = 0.0) (simulated vector fields). **Row 2**: Spatially explicit simulations, strong mixing (*D* = 10.0) (simulated vector fields). **Row 3**: CF approximations (analytical vector fields) *D* = 0.0. In all cases, parameters are: *N* = 9, *κ* = 3, *c*_0_ = 1.0, *c* = 0.3; in the four columns: **A**: *b* = 0.3, *r* = 0.01; **B**: *b* = 0.5, *r* = 0.01; **C**: *b* = 0.65, *r* = 0.2; **D**: *b* = 0.6, *r* = 0.01.

**Figure 5.**
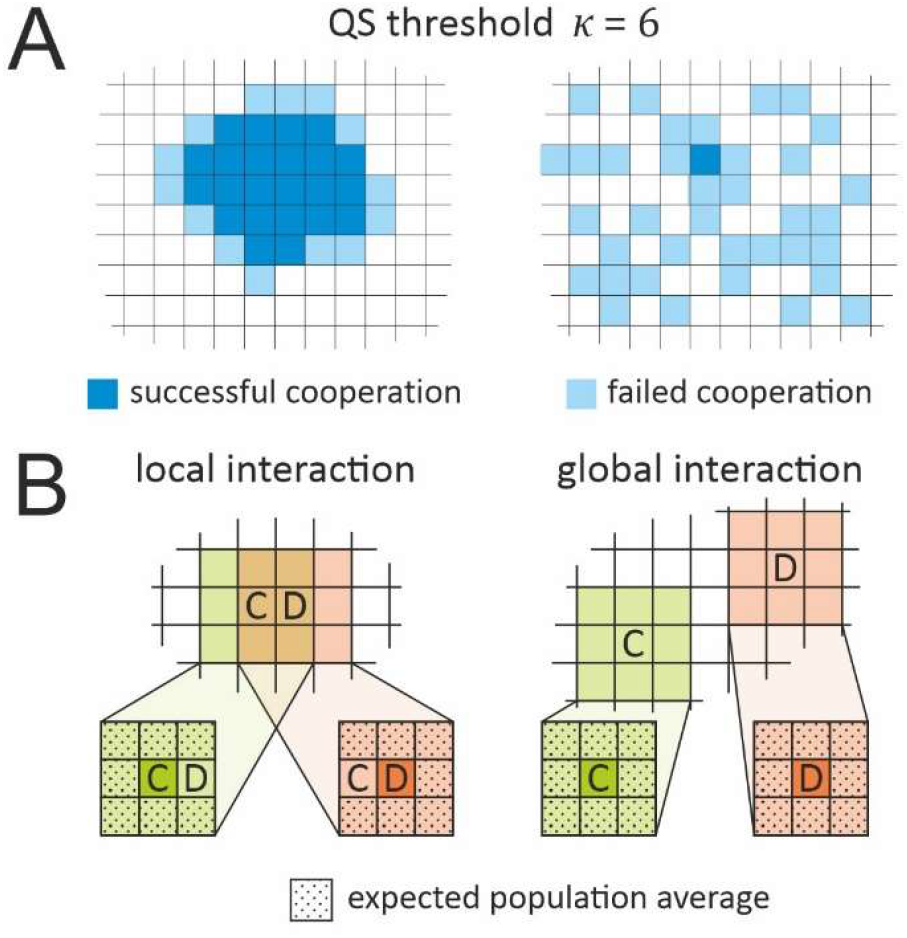
Schematic explanation of the two different effects of spatial constraints in the spatially explicit and implicit models (lattice and CF models, respectively). **A**: The effect of viscosity. Cooperators benefit from slow mixing (viscosity) during the cooperation phase, as their perceived local cooperator density exceeds the population average. Fragmented cooperators fail to achieve the local quorum of cooperation, despite the same global density of cooperators on the lattice. **B**: The effects of local and global competition in well-mixed populations. Local competition between cooperators and parasites (left panel) benefits the parasite because their overlapping cooperation neighborhoods (including themselves) contain an equal expected number of cooperators, but the parasite carries a smaller cost. The cooperator in a distant cooperator-parasite pair (right panel) represents an extra cooperator in its own (otherwise statistically identical) neighborhood compared to its parasitic opponent. This advantage may (over-)compensate its handicap in cooperation cost.

That is, while 1) predicts parasite advantage from intensive mixing of the players on the lattice, in 2) mixing (as represented by competitive interactions occurring between distant individuals) helps cooperation. Separating (in time) the cooperation phase from the competition phase gives an advantage to cooperators in the CF model, which counter-balances the adverse effect of mixing (see the Discussion for more detail on this mechanism). This explains why the viscous (spatially correlated) lattice model and the non-correlated CF approximation often produce surprisingly similar steady state strategy distributions in the parameter space (cf. Figure 6, Appendix 3, Figure S6).

**Figure 6.**
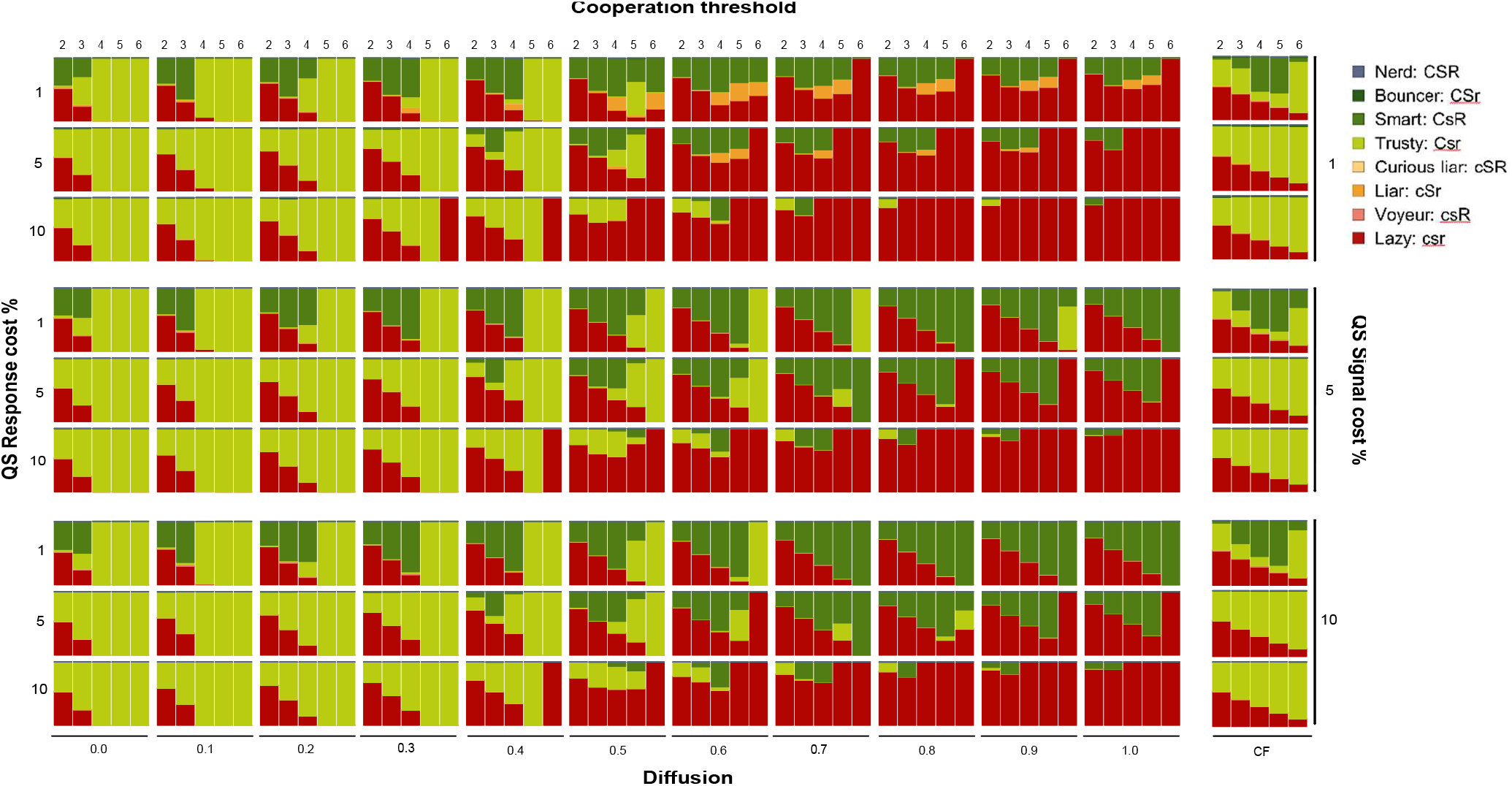
Genotype distributions in steady-state populations of the 8-strategy lattice model and the configuration-field model across the feasible ranges of cooperation threshold (*κ*), QS signal cost (*s*), QS signal response cost (*r*) and agent motility due to diffusion (*D*), with fixed parameters *N* = 9, *c*_0_ = 1.0, *c* = 0.2, *b* = 0.8 in all cases. The functional mutation rate for all strategies is ρ= 10^-4^.

#### 8-strategy model with functional mutations

Extensive simulations on the lattice with all 8 strategies present and mutations allowed (acquiring or losing functional genes for cooperation, QS signal production, or QS signal response) were performed. The parameter space of the model was scanned across four critical parameters: (1) the cooperation threshold *κ* across values from 2 to 6; (2) the QS signal cost *s* with values 1, 5, and 10% of the basic metabolic burden *c*_0_ of the agents; (3) the QS response cost *r* with values 1, 5 and 10% of *c*_0_; and (4) the diffusion (motility) parameter *D* of the agents ranging from 0.0 to 1.0. The genotype distributions at the steady states for all combinations of these parameter settings are shown in Figure 6. The same sets of simulations with different pairs of fixed cooperation cost *c* and cooperation benefit *b* are presented in Appendix 3, Figure S6. The overall trends along the four scanned dimensions of the parameter space are the following.

##### Diffusion (*D*)

The most conspicuous trend is also the most obvious one: increasing agent motility (*D*) benefits the parasitic strategy (Lazy), giving it more access to the (undeserved) cooperation benefit provided by unconditional and conditional cooperators (Trusty and Smart, respectively; Figure 6). There are, however, a few more effects of increasing diffusion which are less obvious. One is the increasing proportion of the Smart (quorum sensing) strategy among the decreasing number of cooperators, which makes perfect sense in a population with local neighborhood configurations highly variable in terms of the number of cooperators in them. It is in this case that the small cost of quorum signal detection pays off by sparing unnecessary cooperation costs at low local cooperator density but switching on cooperation in sufficiently cooperative neighborhoods. At very low and very high agent motilities, however, the local neighborhoods are predictable enough to render QS a futile waste of resources. It is also worth noting that the CF approximation (rightmost columns of panels in Figure 6 and Appendix 3, Figure S6) behaves almost like the lattice model at low mixing (small *D*), once again underlining the negative effect of the overlapping cooperation neighborhoods of adjacent competitors.

##### Cooperation threshold (*κ*)

The quorum signal response threshold (the minimum number of signals in a neighborhood that switches on conditional cooperation in Smart agents) and the cooperation threshold (the minimum number of actual cooperators providing the cooperation benefit for the focal agent of the neighborhood) are assumed to be the same in all our models. For cooperation to be an option, *κ* = 2 is the minimum requirement. The potential maximum number of cooperators within a Moore neighborhood is *κ* = 9, but cooperation would be compulsory for all the agents at *κ* = 9, and we have found it almost impossible to evolve for *κ* > 6, so the feasible range to scan was *κ* = 2 to 6. Figure 6 clearly shows that, contrary to common intuition, cooperation evolves to its highest frequency in the population at higher intermediate values of the cooperation threshold (at *κ* > 3) (Archetti and Scheuring 2010, 2012). Approaching *κ* = 6 unconditional cooperators (Trusty) often completely exclude parasites (Lazy) at low agent motility (*D*), whereas parasite/cooperator coexistence is the generic outcome of spatial simulations at low *κ* and *D*, like in the CF model. Towards the high end of the *κ* scale (upward from *κ* = 6), cooperation abruptly disappears in a phase-transition-like manner, leaving the system in the monomorphic Lazy steady state across the whole parameter space with *κ* ≥ 7 (data not shown).

##### QS response cost (*r*)

The metabolic cost of QS signal response (signal detection and intracellular signal transduction) is assumed to be low compared to the cost of cooperation (*r* ≪ *c*). This is reasonable, given that cheap cooperation coordinated by expensive signal response is certainly a losing strategy against cheap cooperation with no QS at all because constitutive cooperation would then be cheaper than listening to the QS signal. If signal response is cheap (*r* = 1.0), then a considerable fraction of cooperating agents maintain, and use it within almost the entire range of the parameter space, except where the parasitic Lazy population goes extinct. Obviously, with all parasites wiped out, it is not worth listening to QS signals anymore, so Trusty takes over. Also, at low agent motility (smaller *D* values), maintaining a more expensive QS signal response (*r* = 5.0 and 10.0) proves not to be feasible. An interesting effect of cheap to moderately expensive QS signal response (*r* = 1.0 and 5.0) at *D* = 0.4 and *κ* = 6 is that it helps to eliminate the parasite, even though it is ultimately not present in the steady-state population. More expensive signal response (*r* = 10.0, with all other parameters the same) results in parasite takeover.

##### QS signal cost (*s*)

Recall that cooperation always means synchronized signaling in our models, so an agent that issues an extra signal is a liar: it is either not a cooperator but pretends to be one, or is a cooperator, but promises more cooperation than it actually provides. Given that cheating is usually a profitable strategy in cooperative situations, it is quite surprising how efficient QS is in eliminating it. Liars (**cSr** genotypes that do not cooperate and do not listen to QS signals) appear only in a narrow section of the parameter space (at low costs for both QS signaling *s* and QS signal response *r*), and even there they are present at low frequencies. Cooperators trying to exaggerate their cooperative nature (Bouncer: **CSr,** and Nerd: **CSR**) do not appear in the simulations in frequencies above their mutation-selection balance.

## Discussion

### Cheap quorum sensing supports cooperation

Since the functioning cooperation allele **C** is also a cue for potential interacting partners on future cooperation, possession of a fully functional QS system means harboring a functional response gene **R** and a conditionally expressed cooperation allele **C**. Figure 6 reveals that in spatially explicit, viscous systems a cheap response function (small *r*) of QS substantially increases the overall propensity for metabolically much more expensive cooperation. In this case, it is mostly conditional cooperators (Smart, **CsR**) that coexist with defectors (Lazy, **csr**). More expensive **R** alleles prevent both signal response and cooperation, with only defectors surviving at moderate to high cell motility (*D*). In the CF approximation, cheap signal response (*r* = 1) allows the Smart strategy to attain high frequency in the steady state, but it does not noticeably contribute to the evolutionary success of cooperation: at higher response costs the Trusty population performs just as well without the **R** allele (see Figure 3).

### Spatial correlations do not favor cooperation in all contexts

The well-known prediction of almost any game theoretical model on “uninstructed” cooperation (in which individuals have no prior clue on the intent of cooperative or defective behavior of their interacting partners) is that the intensive mixing of cooperators and defectors destroys costly cooperation through the inevitable fitness advantage of defectors in cooperator/defector encounters. This universal conclusion holds true in the threshold public goods game with competition restricted to neighbors, even if all cooperators are assumed to constitutively broadcast their cooperative nature with a signal that their potential partners may detect at a small cost and switch on or off their cooperation mechanism accordingly. This is obvious from Figure 6: defectors take over as the mixing parameter *D* increases in the agent-based, spatially explicit simulations, irrespective of all other parameter values. However, maintaining spatial correlations does not always favor cooperation *in any context*. On the one hand, spatial correlations always favor cooperation during the fitness acquisition phase, as it allows cooperators to be more likely surrounded (and helped) by other cooperators; thus, cooperation is more likely to succeed. On the other hand, spatial correlations favor defectors at the competitive reproduction stage as it hurts successful cooperators to compete against (potentially also successful) defector neighbors. In other words, cooperators benefit from cooperating locally but competing globally for reproduction, suggesting an ambivalent effect of spatial mixing on cooperation in TPGG. This ambivalent effect is dependent on the temporal scales of the two types of interaction assumed. The advantage of cooperators in the CF model (as opposed to that of cheaters in the well-mixed spatially explicit simulations) hinges on the implicit assumption of the instantaneous collection of fitness by all agents in the cooperation phase, and their also instantaneous competitive interaction in the imitation phase with a random opponent *at a later moment in time*. In both models, intensive population mixing is assumed, but in the CF the two interaction phases are farther apart in time, whereas in the spatially explicit simulations they are simultaneous. It should also be noted here that mixing may also have a detrimental effect on competing cooperators by allowing more frequent cooperator-parasite encounters in space. This, and the dissolution of cooperator clumps eventually always lead to parasite takeover at fast population mixing in the spatially explicit lattice model.

### Quorum sensing evolves if it conveys valuable information

QS is efficient at reducing the overall metabolic cost (and thus at increasing the average fitness of the population) if the expected number *E*(*n*_c_) of cooperators per neighborhood is close to *κ* (the cooperation threshold) in the actual steady state of the population, and the variance of the same variable, *V*(*n*_c_) is relatively high. These are the conditions at which it is difficult to predict if a particular neighborhood does or does not have the quorum of cooperators (*n*_C_ ≥ *κ*); therefore, the information that the QS system provides is of the highest fitness advantage. This is quite obvious in the spatially explicit lattice model versions, but also applies to the CF approximation that explicitly considers the stochastic heterogeneity of neighborhood composition, too. Of course, the MF approximation never yields QS because it assumes all neighborhoods to be identical and thus completely predictable, in which case maintaining the signal-detecting **R** allele (i.e., expressing the signal receptor and the signal transduction system) would be a waste of resources and is thus selected against.

### High cooperation thresholds favor monomorphic equilibria

A counter-intuitive prediction of the spatially explicit version of the threshold public goods game model is that the higher the cooperation threshold *κ* (i.e., the more cooperators are needed for harvesting the cooperation benefit), the higher the steady-state frequency of the cooperators will be (Archetti and Scheuring 2010, 2012), and at moderately high *κ* values the defectors may be eradicated altogether. On the one hand, this is a natural consequence of requiring more cooperators per interaction neighborhood which translates to higher overall (regional) cooperator abundance in mixed steady states, but on the other hand, it is surprising that increasing *κ* does not favor defection – on the contrary, it seals the fate of defectors. This effect shows up at low agent mobility in the spatially explicit simulations, but the trend of increasing cooperator frequency with increasing *k* can also be observed in the CF approximation, which does not provide the advantage of cooperator clumping at all, as it assumes complete mixing of the population. At *κ* values too close to the neighborhood size the steady-state populations are also monomorphic, but then the defectors take over the lattice in a phase transition-like manner with increasing *κ*.

### Liars are kept at bay if cooperativity is always visible, even if fake signaling is unconstrained

Yet another interesting prediction of the spatially explicit agent-based model is the limited range of the parameter space in which cheating through issuing fake quorum signal (lying) can evolve at all: the only section of the parameter space of the 8-strategy lattice model in which the Liar strategy shows up is at very low signal cost and low response costs (Figure 6, *s* = 1, *r* = 1 and *r* = 5), and even there it attains almost negligible frequencies, in spite of the fact that the **S** locus is free to mutate back and forth, just as the other two (**C** and **R**). The fact that lying is always disadvantageous from the viewpoint of evolving cooperation can be seen in the reduction of the frequency of cooperative strategies wherever Liars occur in the population, but in all other parts of the parameter space Liars are missing from the steady state. Lying is inefficient due to the fact that cooperators can produce a baseline signal for free in our model while liars have to pay for the same signal. In other words, there is a condition-dependent trade-off that favors honesty. Theoretical models of honest signaling have shown that honesty is maintained by such condition-dependent trade-offs (Hurd 1995, Számadó 1999, Lachmann et al. 2001, Bergstrom et al. 2002, Számadó et al. 2019, 2023), instead of the equilibrium cost of signals (a.k.a. ‘handicaps’) as predicted by the erroneous Handicap Principle (Zahavi 1975) (see (Penn and Számadó 2020) for discussion), and later costly signaling models (Grafen 1990, Godfray 1991). While such trade-offs are difficult to measure, a recent experiment (Számadó et al. 2022) supports the key role of condition-dependent trade-offs in maintaining honesty. In turn, our model gives an example of cheap (cost-free) and honest signaling under conflict of interest. This possibility was predicted long ago (e.g. (Számadó 1999, 2011, Lachmann et al. 2001), but implementations of such systems are few and far between.

All in all, our results demonstrate that cue-based quorum sensing can maintain cooperation at a high level, partly due to the conventional kin selection mechanism of cooperative games, partly as the result of specific spatial effects in local public goods games. The frequency of communication cheaters is constrained in such a system by the condition-dependent trade-off in signal production (i.e., signaling is free for honest signalers but costly for cheaters).

While several models have investigated potentially deceptive QS strategies in bacteria (Czárán and Hoekstra 2009, Wang et al. 2020, Gurney et al. 2020), there are key differences between the current investigation and the previous ones. First of all, we investigate a cue-based system, in which the signalling molecule is the same as the public good. This provides dynamics different from those of the traditional QS systems with separate signal and public good molecules, which, therefore, have to be synthesized along different metabolic pathways. Of course, not all QS systems are cue-based, but our results suggest that whenever it is cue-based, it strongly favors honesty and cooperation. Secondly, like Czárán & Hoekstra (2009), we investigate the full range of 8 strategies possible, given the assumption that each of cooperation, signaling and signal detection may be ON or OFF, but in the present model cooperators that are also signalers are a new type of cheat: those who exaggerate their promise of cooperation. In spite of even more cheating strategies being possible, we find a substantially reduced prevalence of cheats in the population. Last but not least, we have decomposed and separated the effects of spatial and temporal constraints present in the spatiotemporal agent-based simulation model by partially adding or eliminating them in the mean-field and configurational-field approximations. This allowed us to resolve simple scenarios, and the approximations provided benchmarks for the agent-based simulations. While this allows a thorough study of the cue-based QS system, there are several issues that merit further investigation. First, we assumed that the cooperation threshold is the same as the activation threshold for conditional cooperators. The reasoning behind this assumption is that conditional cooperators switching on too early or too late will be selected against. This assumption can be investigated with a slightly modified version of this model. The main reason for not including this quite obvious and reasonable complication in the current version is to keep the complexity of the investigation under control. Second, perhaps more interestingly, our results beg the question: why is it that not all QS systems are cue-based? If cue-based systems promote honesty and cooperation, then one might expect such systems to be widespread in nature. Yet, while there are cue-based systems besides the nisin system in *Lactococcus lactis*, like the mutacin 1140 system in *Streptococcus mutans* (Kreth et al. 2005), the mersacidin system in *Bacillus sp.* (Schmitz et al. 2006) or the listeriolysin S (LLS) system in *Listeria monocytogenes* (Cotter et al. 2005, Freitag et al. 2009), there are many more examples in which the signal molecule is different from the public good. It is conspicuous that most known cue-based QS systems regulate autoinduced bacteriocin (lantibiotic) production, i.e., toxin excretion, possibly deployed against unrelated competitor strains. Perhaps cue-based systems are also constrained by some other, still unknown, genetic or ecological mechanism. This leads to the issue of how cue-based QS systems had evolved in the first place – a potentially rewarding focus for future investigations.

## Methods and models

### Mean-field (MF) model

In Appendix 1, we construct the mean-field version of the model, with the assumptions that population size (*P*), interacting group size (*N*) and the cooperation threshold *κ* are all very large, while *N*⁄*P* ≪ 1 and *κ*/*N* → *K*. In this limit all the dynamical effects are averaged across the entire habitat so that the fitnesses of the strategies depend only on the average frequencies of the strains present. On the basis of simple fitness considerations, it holds that only combinations of the Lazy, Trusty and Smart strategies can constitute any equilibrium state in the MF approximation, thus we have to analyze the dynamics only for these three strategies. To see this, let us assume indirectly that there is a fourth strategy *S* present in the equilibrium. Since at least one of the strategies from among Lazy, Trusty or Smart has a higher fitness than strategy *S* in any actual state of the frequency vector ***X***, *S* can not be present in a dynamical equilibrium. For the mathematical formulation of the MF model see Appendix 1.

### Configuration-field (CF) model

Next, we assume that the population is still very large (practically infinite), but individuals form random interacting groups of finite size *N*. While in the previous section we considered *N* to be so large that each interacting group consists of strategies in exact proportion to their global frequencies in the population, now we assume that *N* is smaller, and thus different interaction groups with different configurations of strategies are formed in an inherently stochastic manner, due to sampling errors. The two players participating in an elementary game step are randomly chosen members of their own interaction groups (both of them of size *N*) that are drawn at random from the population. Note that assuming The overall fitness for each of the eight strategies is calculated as the weighted average of its local fitness in all possible configurations of the interaction group around a focal individual of the given strategy. Based on these fitness formulae it can be shown again that the feasible strategy set consists only of Lazy, Trusty and Smart (for more details, see Appendix 2).

### Agent-based non-spatial model

The agent-based non-spatial model is an accurate computational realization of the CF approximation in all respects, except the population is now finite (population size *P* = 90.000). Two random samples of size *N* (*N* = 9), drawn from the population independently, represent the interacting groups. One focal agent from each group is chosen at random, and these two focal agents play out the imitation game, with the chance of winning for each dependent on the fitness payoff it has realized within its own interacting group. Payoffs were assigned to each of the two players depending on their own cooperation costs and the number of actual cooperators in their own group, which, in turn, depends on the local number of signalers and conditional cooperators within the corresponding interacting group. The updating rule is random, meaning that each update step consists of the choice of a random pair of players and their cooperation groups, and the competitive imitation step between them. One generation comprises *P*/2 such steps so that each agent participates in one update per generation on average.

### Agent-based spatial (lattice) model

The spatially explicit model follows the agent-based algorithm, with the modification that now the agents are arranged in a 300×300 square lattice to implement the spatial constraints of localized interactions and limited agent mobility. The lattice is of toroidal topology (with its opposite edges merged) to avoid edge effects. The players in each updating step are immediate neighbors of each other with their interacting groups consisting of the two overlapping 3×3 sub-lattices centered on them (i.e., the interacting group of a player occupies its Moore neighborhood on the lattice). In an elementary game step, an imitation game is played out between the randomly chosen pair of adjacent agents. Obviously, this means that the interacting neighborhoods of the two players are not independent of one another: the two players are always members of their own, as well as of the other’s, interacting group, and the two interacting groups (neighborhoods) share some other common agents as well. The actual size of the overlapping region depends on whether the players are orthogonal or diagonal neighbors (Figure 5B). Payoffs, and thus also fitness values at interaction, are assigned to each of the two players as in the non-spatial model. For a detailed description and additional information, see Appendix 3.

The limited spatial mobility of the agents on the lattice is scaled by the diffusion parameter *D*, which is the expected number of site swaps between randomly chosen pairs of adjacent agents following each game step. The swapped pairs are chosen independently of the interacting pair.

One generation of the spatial simulation also consists of *P*/2 random elementary interaction steps and the corresponding (random) diffusion steps. Simulations last for *G* = 10.000 generations. Empirical vector fields with particular parameter settings have been produced on the strategy simplex for the lattice model to visualize simulated steady states and trajectories.

## Abbreviations

QS: quorum sensing
MF: mean-field
CF: configuration-field
PGG: public goods game
TPGG: threshold public goods game

## Additional information

### Supplementary information

**Additional file 1: Appendix 1**, Mean-field model, Fig. S1; **Appendix 2**, Configuration-field model, Figs. S2-S5; **Appendix 3**. Individual-based lattice model, Fig. S6.

**Additional file 2**: Wolfram Language code of the configuration-field model to reproduce Figure 3 and Figure 4.

**Additional file 3**: Fortran code of the lattice-based model to reproduce Figure 4.

### Ethics approval and consent to participate

Not applicable.

### Consent for publication

Not applicable.

### Availability of data and materials

Code to reproduce data and figures are available in the **Additional files 2 and 3**.

### Author’s contributions

TC and SS conceived the idea and designed the study, IS and TC analyzed the analytical models, TC designed the individual based model, TC, SS and IZ implemented and run the individual based model and analyzed data, TC, SS, IZ and IS created the figures. All authors contributed to the writing and editing of the manuscript.

## Acknowledgements

None.

