## Appendix 1-3 for "Cue-driven microbial cooperation and communication: evolving quorum sensing with honest signalling"

Tamás Czárán[1], István Scheuiring[1], István Zachár[1,2], Szabolcs Számadó[3,4]

1 Institute of Evolution, Centre for Ecological Research, Konkoly-Thege Miklós út 29-33., H 1121 Budapest, Hungary
2 Department of Plant Systematics, Ecology and Theoretical Biology, Eötvös Lóránd University, H-1117 Budapest, Hungary
3 Department of Sociology and Communication, Budapest University of Technology and Economics, Egry J. u. 1, Budapest, H-1111, Hungary
4 CSS-RECENS Lendület Research Group, Centre for Social Science, Tóth Kálmán u. 4, H-1097, Budapest, Hungary

**This pdf file includes:**
Supplementary text: the mean field model, the configuration field model and additional results for the lattice model
Figs S1 to S6
References

### Appendix 1. The mean-field model

Let $x_1, x_2, x_3, x_4, x_5, x_6, x_7, x_8$ $(\sum x_i = 1)$ denote the frequencies of the strategies of $La, Tr, Bo, Sm$, $Ne, Li, Cl, Vo$, respectively. $x_2+x_3+x_4+x_5 = X_C$ is the total concentration of cooperators, $x_k$ is the cooperation threshold concentration and $X_C + x_3 + x_5 + x_6 + x_7 = X_S$ denotes the total concentration of signals. $N$ is the neighborhood size, which is assumed to be large in this limit. By using the notation introduced in the main text and above the strategies have the following fitness in the mean field limit:

$$W_{La}(X_C) = W_0 - c_0 \left[1 - b\theta(X_C)\right] \tag{1}$$

$$W_{Tr}(X_C) = W_0 - (c_0 + c) \left[1 - b\theta(X_C + 1/N)\right] \tag{2}$$

$$W_{Bo}(X_C) = W_0 - (c_0 + c + s)\left[1 - b\theta(X_C + 1/N)\right] \tag{3}$$

$$\begin{aligned} W_{Sm}(X_C, X_S) = W_0 - (c_0 + r + c)(1-b)\theta(X_C + 1/N) - \\ \left[c_0 + r + c\theta_{S|1-C}(X_C + 1/N, X_S + 1/N)\right]\left[1 - \theta(X_C)\right] \end{aligned} \tag{4}$$

$$\begin{aligned} W_{Ne}(X_C, X_S) = W_0 - (c_0 + r + s + c)(1-b)\theta(X_C + 1/N) - \\ \left[c_0 + r + s + c\theta_{S|1-C}(X_C + 1/N, X_S + 2/N)\right]\left[1 - \theta(X_C + 1/N)\right] \end{aligned} \tag{5}$$

$$W_{Li}(X_C) = W_0 - (c_0 + s)\left(1 - b\theta(X_C)\right), \tag{6}$$

$$W_{Cl}(X_C) = W_0 - (c_0 + s + r)\left(1 - b\theta(X_C)\right), \tag{7}$$

$$W_{Vo}(X_C) = W_0 - (c_0 + r)\left(1 - b\theta(X_C)\right), \tag{8}$$

where

$$\theta(X) = \begin{cases} 1, & \text{if } X \geq x_k \\ 0, & \text{otherwise} \end{cases}$$

,

$$\theta_{S|1-C}(X, Y) = \begin{cases} 1, & \text{if } X < x_k \text{ and } Y \geq x_k \\ 0, & \text{otherwise} \end{cases}$$

are step functions representing the effect of cooperation as functions of the cooperative strategies' concentrations and the signal concentrations. Since $N$ is the neighborhood size, the focal strategy is taken into account with weight $1/N$ in the $\theta$ functions if it cooperates (eqs. 1-5) and/or as a signaler if it signals (eqs. 4,5). The $Ne$ strategy issues an extra signal and it detects signals as well, so this strategy adds with weight $2/N$ to the signal concentration term when we compute $W_{Ne}$ (eq. 5). Naturally, as the the neighborhood size $N$ tends to infinity, $\theta(X_C + 1/N) \to \theta(X_C)$; consequently, the $La$ strategy attains the highest fitness regardless of the frequency distribution of the strategies. That is, $La$ is always the winner of selection in this limit.

If $N$ is finite - even if it is large - and $b$ is sufficiently high compared to the costs $c, s, r$, then besides $La$, the inevitable winner of selection, also the cooperative strategies can be successful. Below we analyse this case. From the definitions of the strategies and the specific assumptions of the mean-field model (i.e., everyone detects the same amount of quorum signal and feels the same effect of cooperation across the habitat) it follows that

$$W_{La} > W_{Li}, W_{Cl}, W_{Vo}$$

and similarly

$$W_{Tr} > W_{Bo}, \quad W_{Sm} > W_{Ne},$$

regardless of the actual strategy distribution. Consequently, we need to consider the relations only among the $La$, $Tr$ and $Sm$ strategies, since all the others are certainly ousted by one of these three. Because all of these strategies are honest, $\theta_{S|1-C}(X, Y) = 0$, thus the fitnesses of these strategies in the mean-field model are simplified to:

$$\begin{aligned} W_{La}(X) &= W_0 - c_0 \left[1 - b\theta(X)\right] \\ W_{Tr}(X) &= W_0 - (c_0 + c) \left[1 - b\theta(X + 1/N)\right] \\ W_{Sm}(X) &= W_0 - \left[c_0 + r + c\right] (1 - b)\theta(X + 1/N) - (c_0 + r)[1 - \theta(X + 1/N)], \end{aligned} \tag{9}$$

where $X$ is the joint concentration of the cooperative $Tr$ and $Sm$ strategies. For sake of simplicity the total concentration of strategies is rescaled to be between zero and 1.

Using standard assumptions of replicator dynamics (Hofbauer and Sigmund 1998) the system is represented by

$$\dot{x} = (W_{La}(X) - \bar{W})x, \tag{10}$$
$$\dot{y} = (W_{Tr}(X) - \bar{W})y,$$
$$\dot{z} = (W_{Sm}(X) - \bar{W})z.$$

where, $x, y, z$ are the frequencies of the feasible $La$, $Tr$ and $Sm$ strategy triplet, and $\bar{W} = W_{La}x + W_{Tr}y + W_{Sm}z$ is the actual average fitness in the population. We studied the qualitative behavior of the above dynamical system by using the the fitness functions of (9).

**Convergence and stability of the $x = 1$ state**

Let us assume that the system is in the $x = 1, y = z = 0$ steady state. We are interested in the stability of this state. Assume that the invader $Tr$ and $Sm$ strategies emerge with frequencies $y$ and $z$ either together or separately. If their total frequency $X = y + z < x_k - 1/N$, then $\theta(X) = \theta(X + 1/N) = 0$ and consequently, $W_{La}(X) > W_{Tr}(X),\ W_{Sm}(x)$, so the invaders couldn't spread. Moreover, it follows from $y + z < x_k - 1/N$ that $x > 1 - x_k + 1/N$, consequently $\dot{x} > 0$ in(10) after invasion, therefore $\lim_{t\to\infty} x = 1$. Thus we have shown that the $x = 1, y = z = 0$ state is an asymptotically stable fixed point of the system if the total initial frequency of invaders satisfies $y + z < x_k - 1/N$.

**Coexistence of $La$ and $Tr$.**

Now let us assume that $X = y + z \geq x_k - 1/N$. Then $\theta(X + 1/N) = 1$ and thus $W_{Tr}(X) > W_{Sm}(X)$, at any parameter setting of the model. That is, the $Sm$ strategy is always beaten by the $Tr$ strategy. Thus we have to consider only the $La/Tr$ subsystem at this initial condition, for which $z = 0$ and $y = 1 - x$ is the frequency of the $Tr$ strategy. As we have shown above if $y < x_k - 1/N$ then $\dot{y} < 0$ in(10), so $Tr$ is selected out. However, if $x_k - 1/N < y \leq x_k$ then $\theta(y) = 0$ while $\theta(y+1/N) = 1$. In this case, the dynamics depend on the parameters of the model. If $b/c > 1/(c_0+c)$, then $W_{La}(y) < W_{Tr}(y)$ and, consequently,

$\dot{y} > 0$, that is, $y$ increases, so after a while $y$ will be greater than $x_k$ and then $\theta(y) = \theta(y + 1/N) = 1$. Thus, $W_{La}(y) > W_{Tr}(y)$, and, consequently, $\dot{y} < 0$, that is, $y$ decreases again. We note here that since $W_{La}(x_k) < W_{Tr}(x_k)$, $x_k$ is not a fixed point of the dynamics by definition, but $W_{La}(x_k + \varepsilon) > W_{Tr}(x_k + \varepsilon)$ for any small $\varepsilon > 0$.). This strange behavior of the system (no fixed point for the replicator dynamics but coexistence anyway) is the consequence that the $\theta$ step function used in the model is nonderivable at the point of the jump. Changing the step function to a continuous sigmoidal benefit function obliterates this singularity by transforming the average benefit of the strategies to a continuous, strictly monotonously increasing function of the strategy distribution. Then the stationary state analyzed above becomes the well known fixed point where $W_{Tr}(y*) = W_{La}(y*)$, with $y* \in (x_k - 1/N \quad x_k)$. Notice that any fluctuation decreasing $y$ below $x_k - 1/N$ drives $Tr$ extinct and fixates $La$ (Fig. S1).

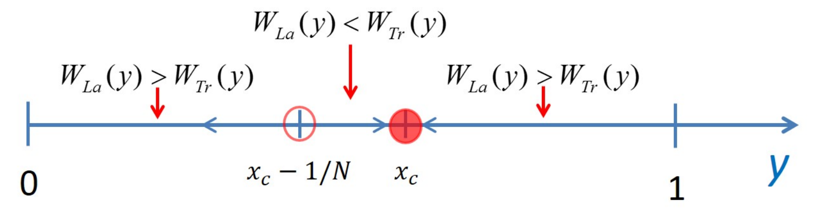


**Figure S1** The qualitative dynamics of the $La/Tr$ system. The $x_k - 1/N$ and $x_k$ points separate three different dynamical behaviours.

As we have shown above the fitness of the $Sm$ strategy is always lower than that of the $Tr$ if $y \geq x_k - 1/N$; therefore, $Sm$ can never invade a $Tr/La$ coexistence.

**Coexistence of $La$ and $Sm$.**

Considering the coexistence of $Sm$ with $La$ the situation is the same as before. The $(z = x_k, x = 1 - x_k)$ is a resting point around which the system fluctuates if $b > (r + c)/(c_0 + r + c)$. However, since $W_{Tr}(X) > W_{Sm}(X)$ if $X = y + z \geq x_k - 1/N$, $Tr$ always invades the $La/Sm$ coalition and excludes $Sm$.

Naturally, if $b/c < 1/(c_0+c)$, then $La$ always has a fitness higher than the alternative strategies $Tr$ and $Sm$, thus $La$ is the only fixed point of the replicator dynamics.

### Appendix 2 The configuration field model

We assume that the interacting group around a focal individual is assembled by drawing $N-1$ individuals at random from the population. The probability of having at least $k$ cooperators (or units of signal) in the group of $N$ individuals thus assembled (including the focal individual) is the weighted average of having $k$, $k$+1,...$N$ cooperators (or signal doses) within the group. Let us denote with $P_l(\vec{x})$ the average probability of having at least $l$ cooperators among the $N-1$ group members if the frequency distribution of the 8 strategies is $(x_1, x_2, ..x_8) = \vec{x}$ in the population. Let $Q_{m|l}(\vec{x})$ be the average probability that the number of signal units is at least $m$ while the number of cooperators is smaller than $l$ in the group of $N-1$ neighbours if the global strategy frequencies are given by $\vec{x}$. Denote the threshold number of cooperators with $k$ ($k\epsilon Z^+, k \leq N$). Using notations and arguments similar to those of the mean-field model we can determine the average fitnesses of the strategies as

$$W_{La}(\vec{x}) = W_0 - c_0[1 - bP_k(\vec{x})] \tag{11}$$

$$W_{Tr}(\vec{x}) = W_0 - (c_0 + c)[1 - bP_{k-1}(\vec{x})] \tag{12}$$

$$W_{Bo}(\vec{x}) = W_0 - (c_0 + s + c)[1 - bP_{k-1}(\vec{x})] \tag{13}$$

$$\begin{aligned} W_{Sm}(\vec{x}) = W_0 - (c_0 + r + c)(1 - b)P_{k-1}(\vec{x}) - \\ (c_0 + r + c)Q_{k-1|k-1}(\vec{x}) - \\ (c_0 + r)[1 - Q_{k-1|k-1}(\vec{x})][1 - P_{k-1}(\vec{x})] \end{aligned} \tag{14}$$

$$W_{Ne}(\vec{x}) = W_0 - (c_0 + r + s + c)(1-b)P_{k-1}(\vec{x}) - \\ (c_0 + r + s + c)Q_{k-2|k-1}(\vec{x})] - \\ (c_0 + r + s)[1 - Q_{k-2|k-1}(\vec{x})][1 - P_{k-1}(\vec{x})] \quad (15)$$

$$W_{Li}(\vec{x}) = W_0 - (c_0 + s)\left[1 - bP_k(\vec{x})\right], \quad (16)$$

$$W_{Cl}(\vec{x}) = W_0 - (c_0 + s + r)\left[1 - bP_k(\vec{x})\right], \quad (17)$$

$$W_{Vo}(\vec{x}) = W_0 - (c_0 + r)\left[1 - bP_k(\vec{x})\right], \quad (18)$$

Like in the the mean field model, it can be shown that $W_{La} > W_{Li}, W_{Cl}, W_{Vo}$ and $W_{Tr} > W_{Bo}$, $W_{Sm} > W_{Ne}$, regardless of the strategy distribution. Thus again only the $La, Tr$ or $Sm$ strategies can be present in any equilibrium state. Assuming random selection of individuals into the interacting groups the fitnesses of these three strategies are

$$\begin{aligned} W_{La}(x,y,z) &= W_0 - c_0[1 - bP_k(x,y,z)] \\ W_{Tr}(x,y,z) &= W_0 - (c_0 + c)[1 - bP_{k-1}(\vec{x})] \\ W_{Sm}(x,y,z) &= W_0 - (c_0 + r + c)(1-b)P_{k-1}(x,y,z) - (c_0 + r)[1 - P_{k-1}(x,y,z)], \end{aligned}$$

with the probabilities computed as

$$P_k(x,y,z) = \sum_{i+j \geq k} f_{i,j}(N-1)x^{N-1-i-j}y^i z^j,$$

$$P_{k-1}(x,y,z) = \sum_{i+j \geq k-1} f_{i,j}(N-1)x^{N-1-i-j}y^i z^j.$$

$f_{i,j}(N-1)$ is the corresponding trinomial coefficient and $x, y, z$ are the frequencies of the $La$, $Tr$ and $Sm$ strategies, respectively. Since there are no non-cooperating signaling strategies in this subsystem, $Q_{k-1|k}(x,y,z) = 0$ at any $x, y, z$ frequency. We consider the same type of replicator equations as above (10) and study this dynamical system qualitatively.

$La$ **is a locally asymptotically stable state of the replicator dynamics.**

We consider the $(x = 1, y = z = 0)$ state and show that the rare invader $Tr$ or $Sm$ strategies have always lower fitness than the resident $La$ strategy. Let us assume first that the invaders follow the $Tr$ strategy with frequency $y << 1$. The success of invasion is determined by the sign of fitness difference of the resident and the invader. After some elementary calculations the fitness difference of $La$ and $Tr$ is determined by

$$W_{La}-W_{Tr} = c[1-bP_{k-1}(1-y,y,0)]+c_0 b[P_k(1-y,y,0)-P_{k-1}(1-y,y,0)].$$

Since $\lim_{y\to 0} P_{k-1}(1-y,y,0) = \lim_{y\to 0} P_k(1-y,y,0) = 0$, the fitness difference tends to $c > 0$ if $y \to 0$. Thus $Sign\{W_{La} - W_{Tr}\} > 0$, that is, the fitness of the invader $Tr$ is always smaller than that of the resident $La$ if the invader is rare. Strategy $Tr$ can not spread if $y << 1$.

Similarly, $La$ is resistant against the invasion of rare $Sm$ $(z << 1)$. The fitness difference of $La$ and $Sm$ is

$$\begin{aligned}(c_0 + r + c)(1-b)P_{k-1}(1-z,0,z)+(c_0 + r)[1 - P_{k-1}(1-z,0,z)]- \\ c_0[1 - bP_k(1-z,0,z)]\end{aligned} \tag{19}$$

which, for the same reason as above, tends to $r > 0$ as $z \to 0$. Strategy $Sm$ can not spread if $z << 1$.

**Neither $Tr$ nor $Sm$ are stable against the invasion of $La$.**

Consider again the fitness differences of $La$ and $Tr$ or $La$ and $Sm$, but at the $y \approx 1$ and $1-y \approx 0$ or at $z \approx 1$ and $1-z \approx 0$ initial values, respectively. Since $P_l(0,1,0) = 1$, for $l \in Z^+$; $l \le N$,

$$\lim_{y\to 1} W_{La}(1-y,y,0) - W_{Tr}(1-y,y,0)\} = c\,[1-b]\,,$$
$$\lim_{z\to 1} W_{La}(1-z,0,z) - W_{Sm}(1-z,0,z) = (r+c)\,[1-b]\,.$$

Since these limits are always positive, $La$ always has higher fitness than $Tr$ or $Sm$ if $La$ is rare, that is, $La$ successfully invades both cooperative strategies.

**The $La$ and $Tr$ can be in stable coexistence.**
We are look for frequencies $y$ where

$$W_{La}(1-y,y,0) = W_{Tr}(1-y,y,0), \tag{20}$$

i.e., where $La$ and $tr$ are at equilibrium. Substituting the fitness values and rearranging (20) yields the condition of fitness equilibrium as

$$\frac{c}{b} = cP_{k-1}(1-y,y,0) + c_0F_k(y) = H_1(y), \tag{21}$$

where $F_k(y) = f_{k-1}(N-1)(1-y)^{N-k}y^{k-1}$. It is easy to see that $H_1(0) = 0$ and $H_1(1) = c$. Since $0 < b < 1$, $H_1(0), H_1(1) < c/b$.

Now we show that $H_1(y)$ has only one maximum ($Max\{H_1(y)\} = H_1(y^*)$) in $y \in [0,\ 1]$. We apply that $F_k(y)$ has a single maximum (see e.g. Archetti and Scheuring 2010) in $y \in [0,\ 1]$. Further, it can be shown that $P_{k-1}(1-y,y,0)$ is a monotonously increasing function with $P'_{k-1}(1,0,0) = P'_{k-1}(0,1,0) = 0$ and with an inflexion point at a $y_k \in (0,\ 1)$. To prove this let us observe that $P_{k-1}(1-y,y,0) = \sum_{i=0}^{N-1}\theta((i+1)/(N-1))f_i(N-1)y^i(1-y)^{N-1-i}$ where $\theta(x)$ is a step function defined as $\theta(x)$=0 if $x < k$ and $\theta(x)$=1 if $x \geq k$. Actually, $P_{k-1}(1-y,y,0)$ is the Bernstein polynomial $B_{N-1}(\theta;y)$ of $\theta(y)$ (Phillips 2003). We introduce the function

$$\sigma_\alpha(y) = \frac{1}{1+e^{-\alpha(y-(k-1)/(N-1))}},\ y \in [0,\ 1], \tag{22}$$

and define

$$T_\alpha(y) = \frac{\sigma_\alpha(y) - \sigma_\alpha(0)}{\sigma_\alpha(1) - \sigma_\alpha(0)}, y \in [0,\ 1]. \tag{23}$$

For every $\varepsilon_1 > 0$ there is an $\alpha_c$ that for every $\alpha > \alpha_c$, $\|\theta(y) - T_\alpha(y)\| < \varepsilon_1$ for every $y \in [0,\ 1]$. That is, $T_\alpha(y)$ is an arbitrarily good continuous approximation of the step function $\theta(y)$. Since the Bernstein operator is monotonous (Phillips 2003),

$$\|B_{N-1}(\theta; y) - B_{N-1}(T_\alpha; y)\| < \varepsilon_2 \tag{24}$$

remains valid for every $\varepsilon_2 > 0$ if $\alpha$ is sufficiently large. $B_{N-1}(T_\alpha; y)$ is an arbitrary good approximation of $P_{k-1}(y, 1-y, 0)$. $T_\alpha(y)$ is a continuous and differentiable function, thus if it is monotonous then its Bernstein polynomial is a monotonous function as well (Phillips 2003). Consequently, $B_{N-1}(T_\alpha; y)$ is monotonously increasing, and because of (24) $P_{k-1}(y, 1-y, 0)$ is a monotonously increasing function as well. Further, the Bernstien operator does not change the sign of the derivatives of a function. That is, since $T'_\alpha(y) > 0$, it means that $B'_{N-1}(T_\alpha; y) > 0$, and consequently $P'_{k-1}(1-y, y, 0) > 0$. Similarly, since the sign of $T''_\alpha(y)$ changes from positive to negative at the inflexion point, so do $B''_{N-1}(T_\alpha; y)$ and $P''_{k-1}(1-y, y, 0)$ (Phillips 2003).

$H_1(y)$ has maximum (or minimum) points where $H'_1(y) = 0$, that is, where

$$cP'_{k-1}(1-y, y, 0) = -c_0 F'_k(y). \tag{25}$$

It is easy to show that $F_k(y)$ is a single-humped function with a maximum point $\hat{y} \in (0,\ 1)$. Similarly, it can be shown that $P'_{k-1}(1-y, y, 0)$ is also a single-humped function with one maximum value in the $[0,\ 1]$ range. Since $F'_k(y) > 0$ in $y \in [0, \hat{y})$, and $F_k(0) = P_{k-1}(1, 0, 0) = 0$, eq. (25) has exactly one solution if

$$\lim_{y\to 1} cP'_{k-1}(1-y, y, 0) < \lim_{y\to 1} -c_0 F'_k(y). \tag{26}$$

Figure (S2) visualizes these two functions by a numerical example. After derivation of the functions and rearranging the above relation we get

$$\lim_{y\to 1} c \sum_{i\geq k} f_i(N-1) \frac{y^{i-1}}{y^{k-2}} \frac{(1-y)^{N-2-i}}{(1-y)^{N-k-1}} (i - (N-1)y) <$$
$$\lim_{y\to 1} -(c_0 + c) f_{k-1}(N-1) \left((k-1) - (N-1)y\right).$$

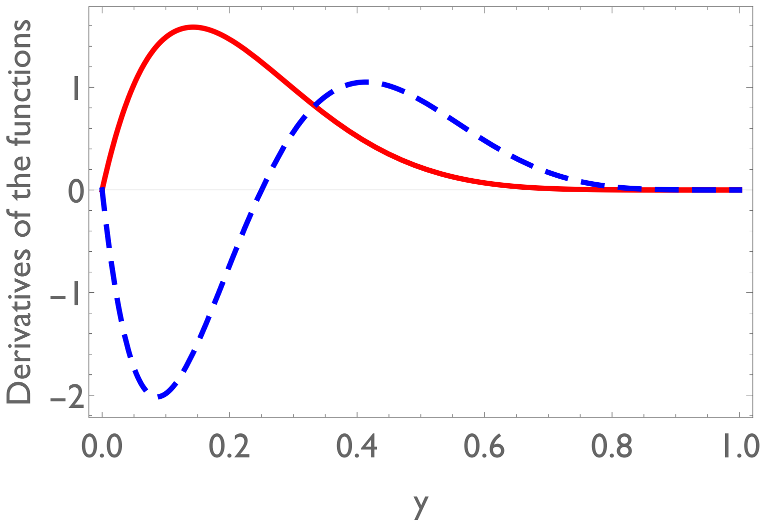


**Figure S2** The shape of $cP'_{k-1}(1-y,y,0)$ (red) and $-c_0F'_k(y)$ (dashed, blue) functions if $H_1(y)$ has one maximum. The intersection of these functions determines the maximum point of $H_1(y)$. $c = 0.5$, $c_0 = 1$, $k = 3$, $N = 9$, in this example.

The left hand side of the relation tends to $-\infty$ and the right hand side tends to $(N-k-1)(c+c_0) > 0$, so relation (26) is satisfied.

Thus we have proven that $H_1(y)$ has only one maximum at $y^*$ in $y \in [0 \quad 1]$. If $H_1(y^*) > c/b$ then there are two solutions $y_u < y^* < y_s$ of eq. (21), where $y_u$ is the unstable and $y_s$ is the stable equilibrium of the $La/Tr$ coexistence. (The stability can be studied by computing the signs of the fitness differences above and below the equilibrium $y_u$ and $y_s$ frequencies). Contrary, if $H_1(y^*) < c/b$ then $W_{La} > W_{Tr}$ for every $y \in [0,1]$, therefore the coexistence of these strategies is not possible (Archetti and Scheuring 2010). In summary, there is at most one stable polymorphic equilibrium $(y_s)$ of $La$ and $Tr$ strategies, and an unstable one $(y_u)$ that separates the trajectories. If $y < y_u$ initially then the system evolves to the pure $La$ strategy $(y = 0)$, while if $y > y_u$ then it converges to the state of coexistence $(y_s)$.

It follows from (21) that there is a critical $b^*$ below which $H_1(y^*) < c/b^*$ for every $c$ and $c_0$, that is, when $La$ always outcompetes $Tr$, and thus the $La$ strategy is the only stable state of the dynamics.

**The stable coexistence of $La$ and $Sm$.**

We perform essentially the same analysis as before; however, in this case equation (21) is modified to

$$\frac{r}{b} = (r + c - c/b)P_{k-1}(1 - z, 0, z) + c_0 F_k(z) = H_2(z) \tag{27}$$

as the condition for the coexistence of $La$ and $Sm$. As before, it can be shown easily that $H_2(0) = 0$ and $H_2(1) = r + c - c/b$, which are both smaller than $r/b$ if $0 < b < 1$. Since $H_2(z)$ is the sum of the same polynomials as $H_1(y)$, we can use the same method as before to show that the $H_2'(z) = 0$ equation has at most one solution in $z \in (0,\ 1)$, and consequently, $H_2(z) = \frac{r}{b}$ has two solutions: $z_u$ and $z_s$ if $Max\{H_2(z)\} > \frac{r}{b}$ in $(0,\ 1)$. There is no solution if $Max\{H_2(z)\} < \frac{r}{b}$. (In the marginal case when $Max\{H_2(z)\} = \frac{r}{b}$, $z_u = z_s$.) The only difference is that $\lim_{z\to 0} H_2'(z)$ can be negative because $r + c - c/b$ can be negative as well.

The following cases are possible:

- If $\lim_{z\to 0} H_2'(z) > 0$ then the situation is the same as before, so there is only one $z^* \in (0,\ 1)$ where $H_2'(z^*) = 0$. This is the point where $H_2(z)$ is maximal (Fig. S3). If $H_2(z^*) > r/b$, then there is an unstable and a stable fixed point of the replicator dynamics of $La$ and $Sm$ in $z \in (0,\ 1)$. Contrary, if $H_2(z^*) \leq r/b$, then $W_{La} > W_{Sm}$ for every $z \in (0,\ 1)$, and thus $La$ beats $Sm$.
- If $\lim_{z\to 0} H_2'(z) < 0$ then, since $P_{k-1}'(1 - z, 0, z) \geq 0$ and $F_k'(z)$ changes the sign exactly once, there are two possibilities: either $H_2'(z) < 0$ for every $z \in [0,\ 1]$ or $H_2'(z)$ changes sign two times in $z \in [0,\ 1]$ (Fig. S4). In the first case $W_{La} > W_{Sm}$ for every $z \in [0,\ 1]$, so $La$ wins the selection (Fig. S5). In the second case $H_2(z)$ has a minimum value at $z^{**}$ and a maximum value at $z^*$. Again if $H_2(z^*) > r/b$, then there is an unstable ($z_u$) and a stable ($z_s$) fixed point of the dynamics of the $La$ and $Sm$ subsystem, otherwise $La$ wins the competition.

**Invasion of two coexistent strategies by the third one**

Let us assume that there is a stable polymorph steady state of $La$ and

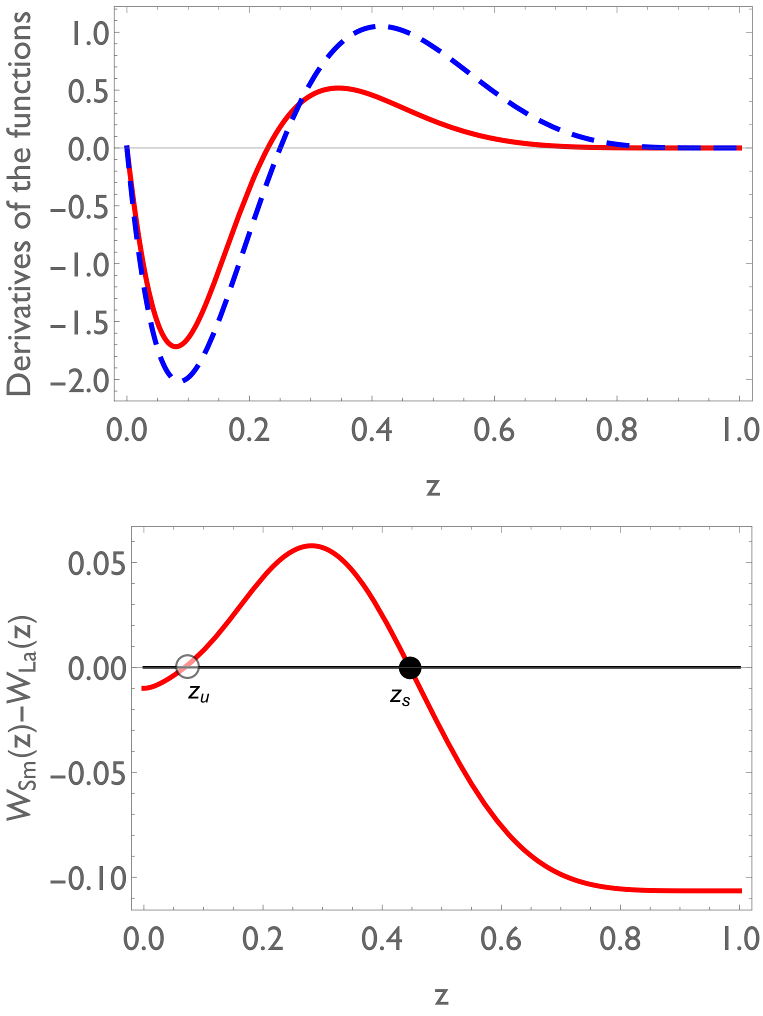


**Figure S3** A case when $H_2'(z)$ is zero at a single $z_{max} \in [0,\ 1]$. **up:** $(r + c - c/b)\, P'_{K-1}(1 - z, 0, z)$ (red), ($-c_0 F'_k(z)$ dashed, blue). **bottom:** The fitness difference of $Sm$ and $La$ in function of $z$. The dynamics has an unstable ($z_u$) and a stable ($z_s$) fixed point. $N = 9, k = 3, c_0 = 1, c = 0.7, b = 0.85, r = 0.01$

$Tr$. The rare $Sm$ can invade this state if its fitness is higher than the fitness of $La$ or $Tr$ in the equilibrium. Let $y_s$ denote the frequency of $Tr$ when $Tr$ and $La$ are in stable equilibrium; thus the fitness of the rare invading $Sm$ is

$$W_0-(c_0+r+c)[1-b]P_{k-1}(1-y_s,y_s,0)]-(c_0+r)[1-P_{K-1}(1-y_s,y_s,0)].$$

Since $P_l(1-y,y,0) = P_l(1-z,0,z)$ if $z = y$, for every $l = 1, 2, N-1$ the above fitness function can be rewritten as $W_0 - [c_0 + r + c][1 -$

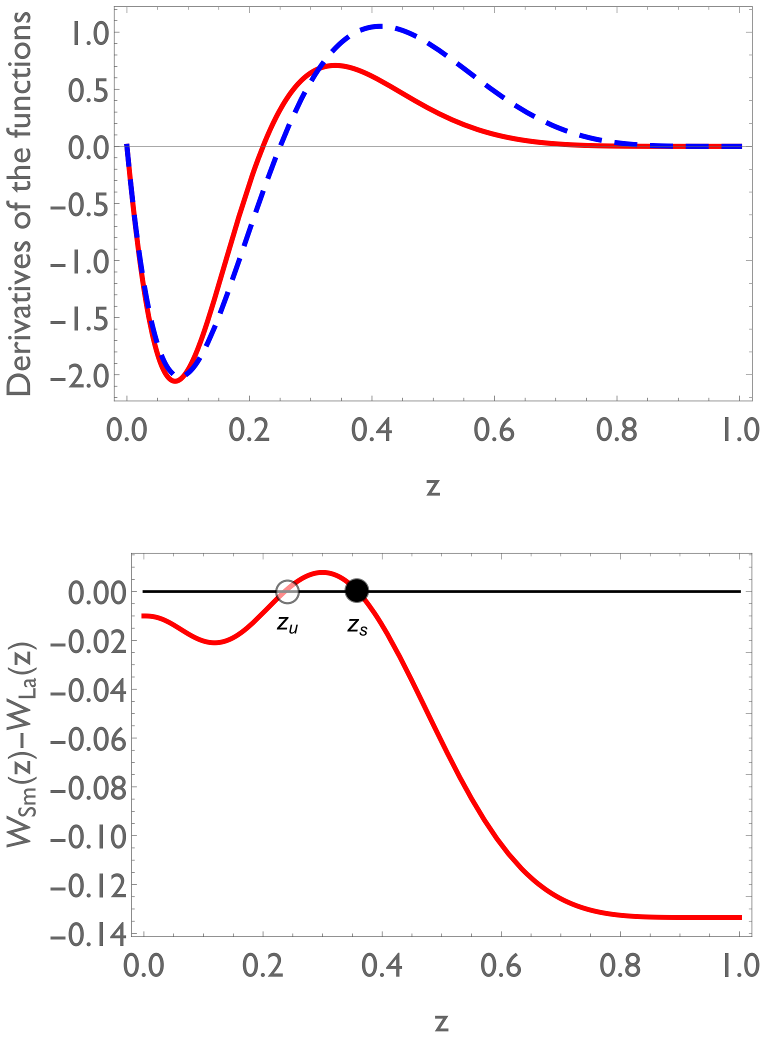


**Figure S4** A case when $H_2'(z)$ becomes zero two times; i.e., there is a local minimum and maximum point of this function.**up:** $(r+c-c/b)\,P_{k-1}'(1-z,0,z)$ (red) -$c_0 F_k'(z)$ (dashed, blue). **bottom:** The fitness differences in function of $z$. Again there are $z_u$ and $z_s$ unstable and stable fixed points of the dynamics. $N=9, k=3, c_0=1, c=0.88, b=0.85, r=0.01$

$b]P_{k-1}(1-y_s,0,y_s)-[c_0+r][1-P_{k-1}(1-y_s,0,y_s)]$ as the fitness of the rare invading $Sm$ at the stable polymorph state of $Tr$ and $La$ . $Sm$ can invade this polymorph $Tr$, $La$ state only if there is a $z_s$ frequency at which $La$ and $Sm$ are in stable coexistence, otherwise the fitness of $Sm$ is always smaller than the fitness of $La$. Let $z_u$ and $z_s$ be the frequency of $Sm$ where $La$ and $Sm$ are in unstable and stable equilibrium. Assume that $y_s \in (z_u,\ z_s)$. Since $W_{Sm}(z) > W_{La}(z)$ for every $z \in (z_u,\ z_s)$, $W_{Sm}(y_s) > W_{La}(y_s) = W_{Tr}(y_s)$, that is, $Sm$ can

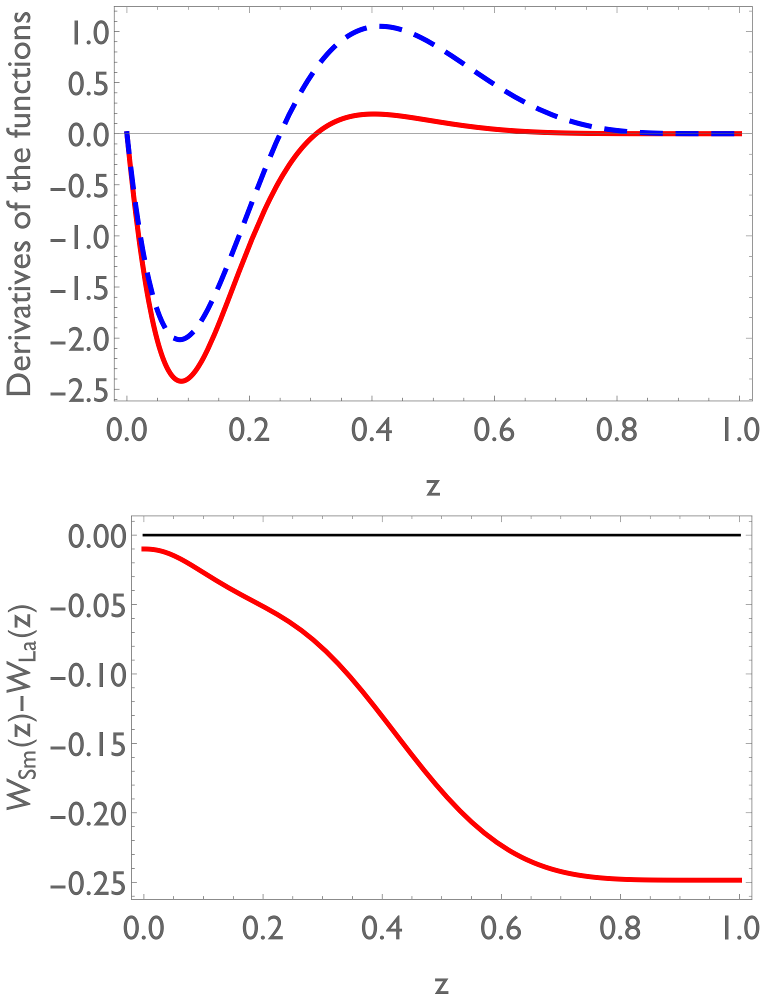


**Figure S5** A case when $H_2'(z) < 0$ and thus $La$ wins over $Sm$. **up:** $(r + c - c/b)\, P_{k-1}'(1 - z, 0, z)$ (red) $-c_0 F_k'(z)$ (dashed, blue). **bottom:** The fitness differences in function of $z$. $N = 9, k = 3, c_0 = 1, c = 0.7, b = 0.65, r = 0.01$

spread when rare in the polymorph state of $La$ and $Tr$. Since $y_s < z_s$, $W_{Tr}(z_s) < W_{La}(z_s) = W_{Sm}(z_s)$ follows; that is, $Tr$ can't spread at the $Sm$-$La$ equilibrium.

Using similar arguments it can be shown that if $z_s \in (y_u,\ y_s)$ the stable polymorph state of $Tr$ and $La$ resists the invasion of the rare $Sm$ strategy, and the polymorph state of $Sm$ and $La$ is unstable against the invasion of $Tr$.

The remaining possibilities are that either $z_s < y_u$ or $y_s < z_u$. Then the system is bistable: neither $Tr$ nor $Sm$ can invade when they are rare. While we couldn't exclude this case per se, we didn't find any parameter combination realizing this situation.

**Coexistence of $Sm$ and $Tr$ is not possible.**

The condition for coexistence is that

$$\frac{c-r}{c-rb} = P_{k-1}(0, y, 1-y). \tag{28}$$

Since $P_k(0, y, 1-y) = 1$ for every $y \in [0,\ 1]$ and the left-hand side is smaller than 1, the above equation can not be satisfied, except in the trivial case of $r = 0$. Otherwise, $Sm$ has a smaller fitness than $Tr$ so it is competed out.

**Coexistence of the three strategies is not possible.**

For the coexistence of $La$, $Tr$ and $Sm$ it is necessary that there is a $(\hat{x} > 0, \hat{y} > 0, \hat{z} > 0)$ where $W_{Tr} = W_{Sm} = W_{La}$. $W_{Tr}$ and $W_{Sm}$ are equal if

$$(c-r)/(c-rb) = P_{k-1}(\hat{x}, \hat{y}, \hat{z}) \tag{29}$$

where $\hat{z} = 1 - \hat{x} - \hat{y}$.

It is clear again, that (29) can be valid only if $r = 0$ and if $\hat{x} = 0$, which excludes the possibility of coexistence of the all three strategies.

### Appendix 3. Lattice model

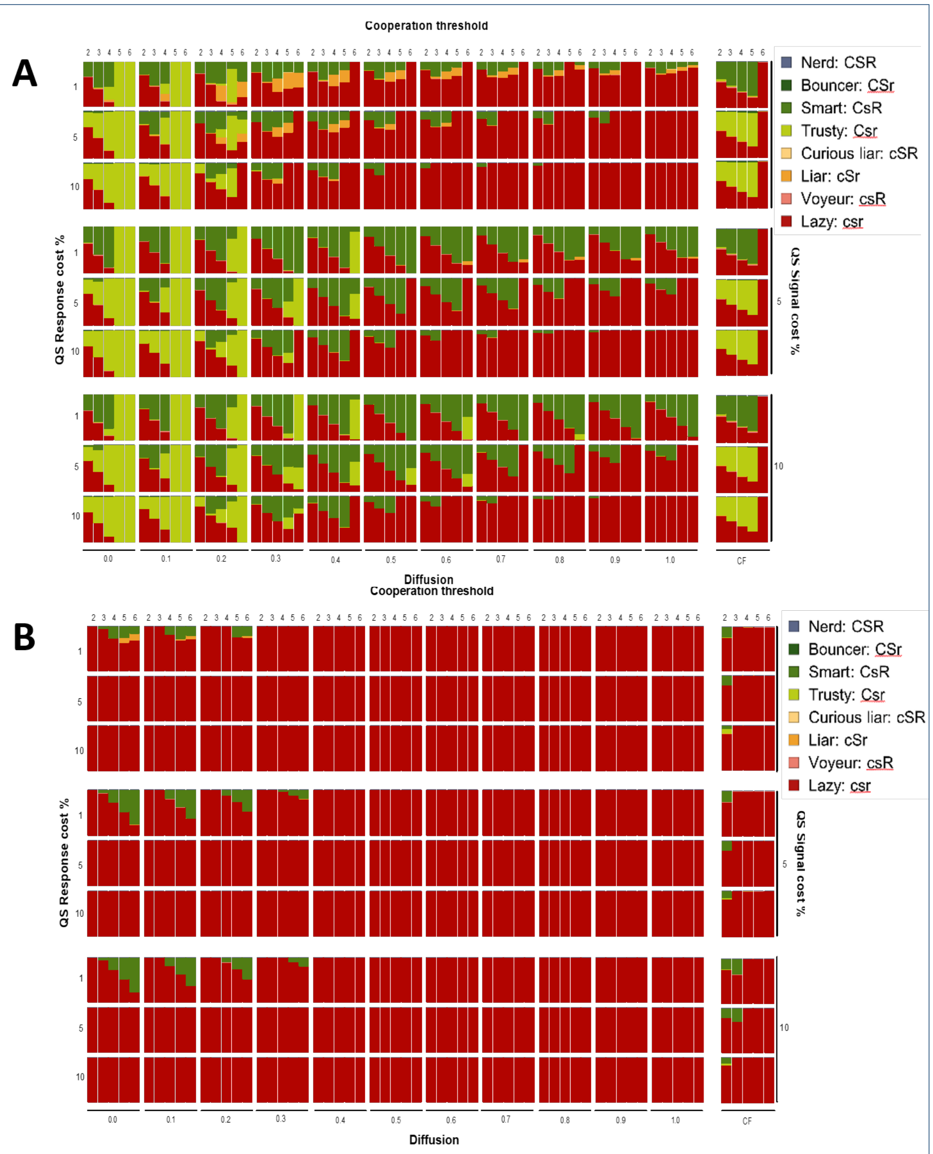


**Figure S6** Genotype distributions in steady-state populations of the 8-strategy lattice model and the CF model across the feasible ranges of cooperation threshold ($\kappa$), QS signal cost ($s$), QS signal response cost ($r$) and agent mobility due to diffusion ($D$), with fixed parameters $N = 9, c_0 = 1.0$ in all cases. The functional mutation rate for all strategies is $\rho = 10^{-4}$. A: $c = 0.3, b = 0.8$; B: $c = 0.3, b = 0.5$.
